# Immune perturbations in human pancreas lymphatic tissues prior to and after type 1 diabetes onset

**DOI:** 10.1101/2024.04.23.590798

**Authors:** Gregory J. Golden, Vincent H. Wu, Jacob T. Hamilton, Kevin R. Amses, Melanie R. Shapiro, Alberto Sada Japp, Chengyang Liu, Maria Betina Pampena, Leticia Kuri-Cervantes, James J. Knox, Jay S. Gardner, HPAP Consortium, Mark A. Atkinson, Todd M. Brusko, Eline T. Luning Prak, Klaus H. Kaestner, Ali Naji, Michael R. Betts

## Abstract

Autoimmune destruction of pancreatic β cells results in type 1 diabetes (T1D), with pancreatic immune infiltrate representing a key feature in this process. Studies of human T1D immunobiology have predominantly focused on circulating immune cells in the blood, while mouse models suggest diabetogenic lymphocytes primarily reside in pancreas-draining lymph nodes (pLN). A comprehensive study of immune cells in human T1D was conducted using pancreas draining lymphatic tissues, including pLN and mesenteric lymph nodes, and the spleen from non-diabetic control, β cell autoantibody positive non-diabetic (AAb+), and T1D organ donors using complementary approaches of high parameter flow cytometry and CITEseq. Immune perturbations suggestive of a proinflammatory environment were specific for T1D pLN and AAb+ pLN. In addition, certain immune populations correlated with high T1D genetic risk independent of disease state. These datasets form an extensive resource for profiling human lymphatic tissue immune cells in the context of autoimmunity and T1D.

## Introduction

Type 1 diabetes (T1D) results in life long exogenous insulin dependence following autoimmune destruction of pancreatic islet β cells^1^. While β cell autoimmunity may start beforehand as suggested by the appearance of β cell-specific autoantibodies (AAb), the disease can take years to progress to overt hyperglycemia^2^. Evidence of β cell mass and insulin secretion can be detected even in long-standing T1D patients^3,4^, with clinically beneficial effects associated with residual β cell function^5^. T1D genetic risk is primarily associated with human leukocyte antigen (HLA) loci but also with genes involved in regulating immune responses^2,6–8^. Accordingly, T cell anergy-inducing immunotherapy shows efficacy in individuals at risk for imminent T1D onset but only delays T1D onset by ∼24 months^9,10^. The development of therapies capable of halting autoimmune progression and preserving β cell function requires further understanding of the complex immune cell responses that occur pre- and post-T1D diagnosis.

Human immunophenotypes associated with T1D include T cell^11–13^, B cell^14,15^, and natural killer (NK) cell^16^ perturbations as well as accelerated immunological aging^17^ in peripheral blood mononuclear cells (PBMCs). While surveying PBMCs can be informative, the immune system operates primarily within tissue microenvironments and disease-specific signatures may not be fully recapitulated in circulating immune cells ^18–22^. Indeed, pancreatic islets of T1D individuals contain β cell antigen-specific CD8+ T cells^23^, and the frequency of CD8+ T cells that recognize β cell antigens in the pancreas, not the blood, distinguishes T1D from healthy donors^24^. Surveying immunophenotypes in the pancreas itself has greatly increased our understanding of T1D pathogenesis^25–30^; however, the immunological processes that occur in pancreas-draining lymphoid tissues that may initiate or exacerbate T1D development and progression remain largely unclear.

Evidence suggests that pancreatic lymph nodes (pLNs) harbor immune cells that participate in T1D autoimmunity^31–34^. Studies in the non-obese diabetic (NOD) mouse model of T1D suggest that pLN lymphadenectomy early in autoimmune development protects against diabetes^31^. Further, pLNs in the mouse model contain stem-like β cell-specific memory CD8+ T cells, which seed cytotoxic and terminally-differentiated effector cells that traffic to the pancreas and eliminate β cells^34^. In humans, pLN from T1D donors contain insulin peptide-reactive T cells^32^, and are enriched for β cell antigen-reactive T cell receptor (TCR) clones^33^. pLN from T1D donors also contain phenotypic alterations that suggest a more proinflammatory T cell population, specifically increased frequencies of Th17 effector CD4+ T cells with reciprocal decreases in regulatory T cell (Treg) and follicular regulatory T cell frequencies^35,36^. While these observations implicate the pLN as a hub of T1D autoimmunity, due to the rarity of these sample types, there has not been a comprehensive unbiased analysis of human pLN that effectively captures this immune environment.

To address these knowledge gaps, we profiled lymphoid immune perturbations across T1D autoimmune development within pLNs, mesenteric lymph nodes (mLNs), and spleen from control non-diabetic autoantibody negative (ND), non-diabetic AAb+, and T1D donors. We performed deep immunophenotyping of these tissues using the complementary methods of high-parameter flow cytometry (n = 46 donors) and cellular indexing of transcriptomes and epitopes by sequencing (CITEseq; n = 18 donors). Taken together, our findings reveal novel immunophenotypes in pLN and mLN from both AAb+ and T1D donors that provide new insights into T1D pathogenesis.

## Results

### Immune profiling of pancreatic, mesenteric, and splenic lymphatic tissues

Through the Human Pancreas Analysis Program (HPAP)^37^, we collected ND, AAb+, and T1D donor spleens and lymph nodes that drain the pancreas, including pLNs (pancreatic head, body, and tail) and mLNs (superior mesenteric artery termed SMA mLN, and upper mesentery termed MES mLN). Donors were age, race, and sex matched across the disease groups (Extended Data Tables 1-2), with tissue samples being processed and analyzed at the same facility over the course of the study. T1D donors were diabetic for around 6 years on average, with only two donors having diabetes >8 years. We used two complementary techniques for deep immune cell profiling of donor tissues: high-parameter flow cytometry on 256 unique samples (18 ND, 10 AAb+, and 18 T1D donors, Extended Data Table 1) and CITEseq single cell analysis on 650,085 cells post quality control (6 ND, 5 AAb+, and 7 T1D donors, Extended Data Table 2). All data are publicly available on PANC-DB (https://hpap.pmacs.upenn.edu/) or Genbank (GSE221787).

We first used a high parameter flow cytometry panel to assess inter-tissue immune differences amongst all major immune cell populations across several immune lineages, including T cells, B cells, innate cells, and subsets thereof (Figure 1A-C, S1). The spleen showed a unique immune profile compared to LNs (Figure 1A-E), with an increased frequency of NK cells (Extended Data Fig. 2A), monocytes, and CCR7-T cell subsets including effector memory (Tem) and Tem CD45RA+ (Temra) T cells (Figure 1C,E). Some subsets were evenly distributed across the spleen and LNs, including B cells (Figure 1C, S2B). In contrast, the LNs were enriched for T cells, particularly CD4+ T cells and naive T cell subsets (Figure 1C-E, S2C-D). The pLNs and mLNs were highly similar in immune cell composition, with few exceptions (Extended Data Fig. 2E-I). Innate lymphoid cells (ILCs) were slightly decreased in frequency within the MES mLN (Extended Data Fig. 2E), CD8+ central memory T cells (Tcm) had a slightly higher frequency occurring in the pLN-Tail (Extended Data Fig. 2F), and NK cells had a slightly decreased frequency in the pLN-Tail (Extended Data Fig. 2A).

**Figure 1.**
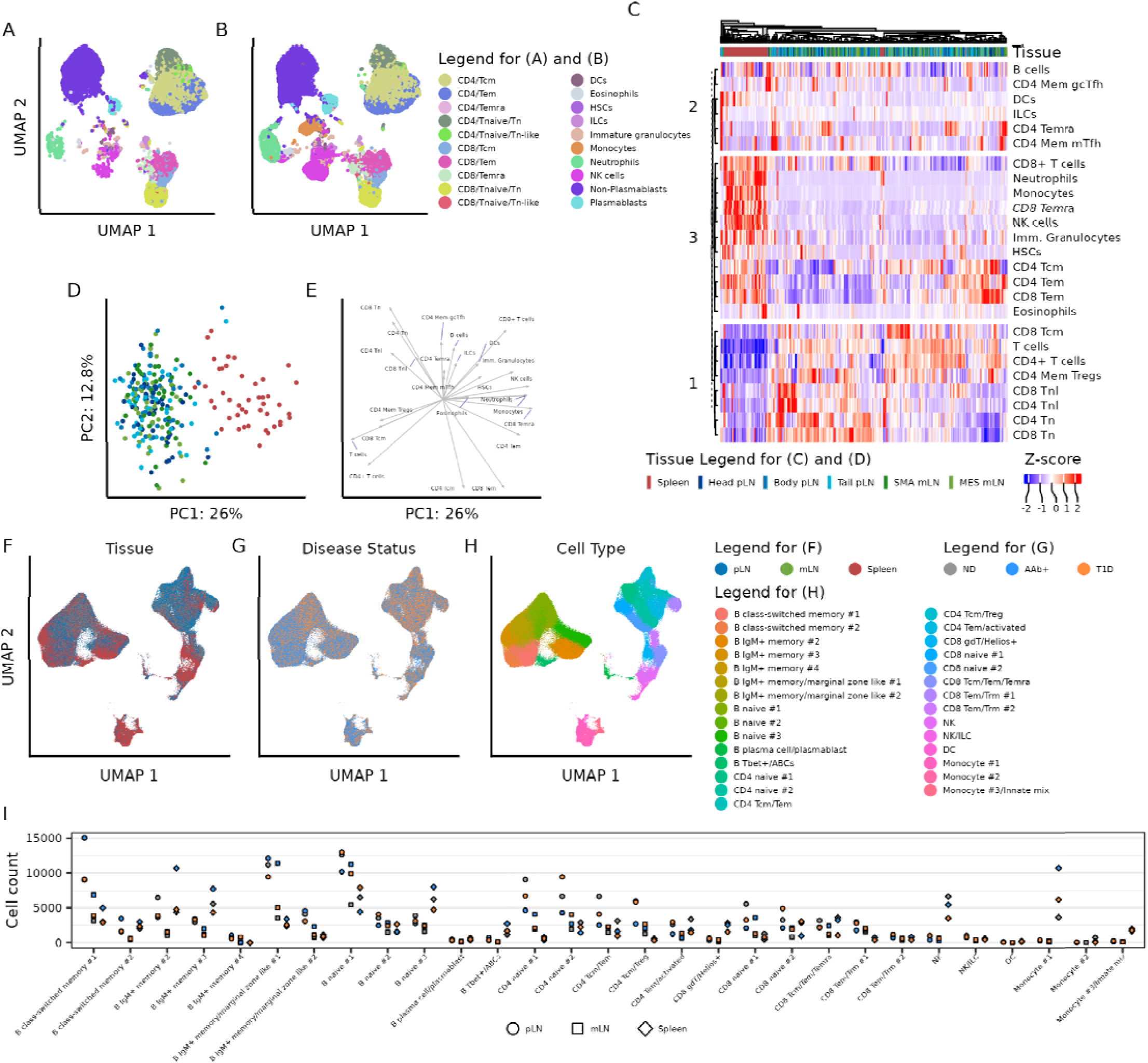
Immune profiling of pancreatic, mesenteric, and splenic lymphatic tissues. (A) UMAP representation of pLN-Tail lymphocytes and (B) splenocytes, highlighting immune lineage populations detected with flow cytometry. (C) Hierarchical clustering of all samples within the flow cytometry dataset using major immune lineage populations. Coloring of the heatmap represents an individual sample’s Z-score within the respective immune population. Immune lineage population clustering patterns were labeled and partitioned at k-means clustering level 3. (D) PCA and (E) biplot of all flow cytometry samples using frequencies of major immune lineage populations. (F) UMAP representation of the RNA component of the CITEseq dataset colored by tissue origin, (G) disease status, and (H) cluster annotation of each cell. (I) Number of cells within each cluster, further subsetted by tissue type and disease state.

In parallel, we performed CITEseq on a subgroup of the same donors, allowing for deep resolution of immune subsets in pLN, mLN, and spleen samples (Figure 1F). No specific immune population selectively resolved by disease state (Figure 1G). Major clusters identified include B cells, T cells, and innate immune cells (Figure 1H), with surface epitopes (Extended Data Fig. 3A) and gene transcripts (Extended Data Fig. 3B) defining subsets thereof. Some clusters in B and T cells contain over 30,000 cells, with at least a few hundred cells in less abundant populations, allowing for deep characterization of rare immune cell populations of interest (Figure 1I). The spleen contained the majority of innate immune cells, while the pLN and mLN samples contained the majority of T cells (Figure 1F,H,I), closely resembling the flow cytometry dataset.

### AAb+ and T1D associated shifts in immune phenotypes

To define immune cell composition and phenotypic changes in T1D development, we compared immune cell population frequencies between ND, AAb+, and T1D donors. As immune cell subsets did not differ in frequency between LN draining the pancreas head, body, and tail or the SMA or MES LNs (Extended Data Fig. 2), we binned different LNs by anatomical origin, either pLN or mLN, for subsequent analysis. Using hierarchical clustering, we observed three different groups of disease-associated immune cell signatures, particularly in the pLN and mLN (Figure 2A). Group 1 immune populations generally decreased in frequency in AAb+ and further in T1D donors, such as CD4+ naive T cells (Tn) cells and CD25+ CD4+ Tem/Temra within the pLN and mLN. Group 2 contained pLN and mLN immune populations, including CD25+ and CD38+ T cell subsets, that decreased in frequency in AAb+ compared to ND donors and maintained a lower frequency or rebounded to normal levels in T1D. Group 3 populations increased in frequency across disease status or were elevated only in T1D. Lymphocyte subsets following this pattern included pLN and mLN populations expressing the IL7 receptor (CD127), the putative residency marker CD69, memory CD27+ B cells, and cytotoxic CD56_dim_CD16+ NK cells. Overall, immune cell phenotypes are modulated in the pLN and mLN of AAb+ and T1D donors, with changes predominantly occurring in the pLN.

**Figure 2.**
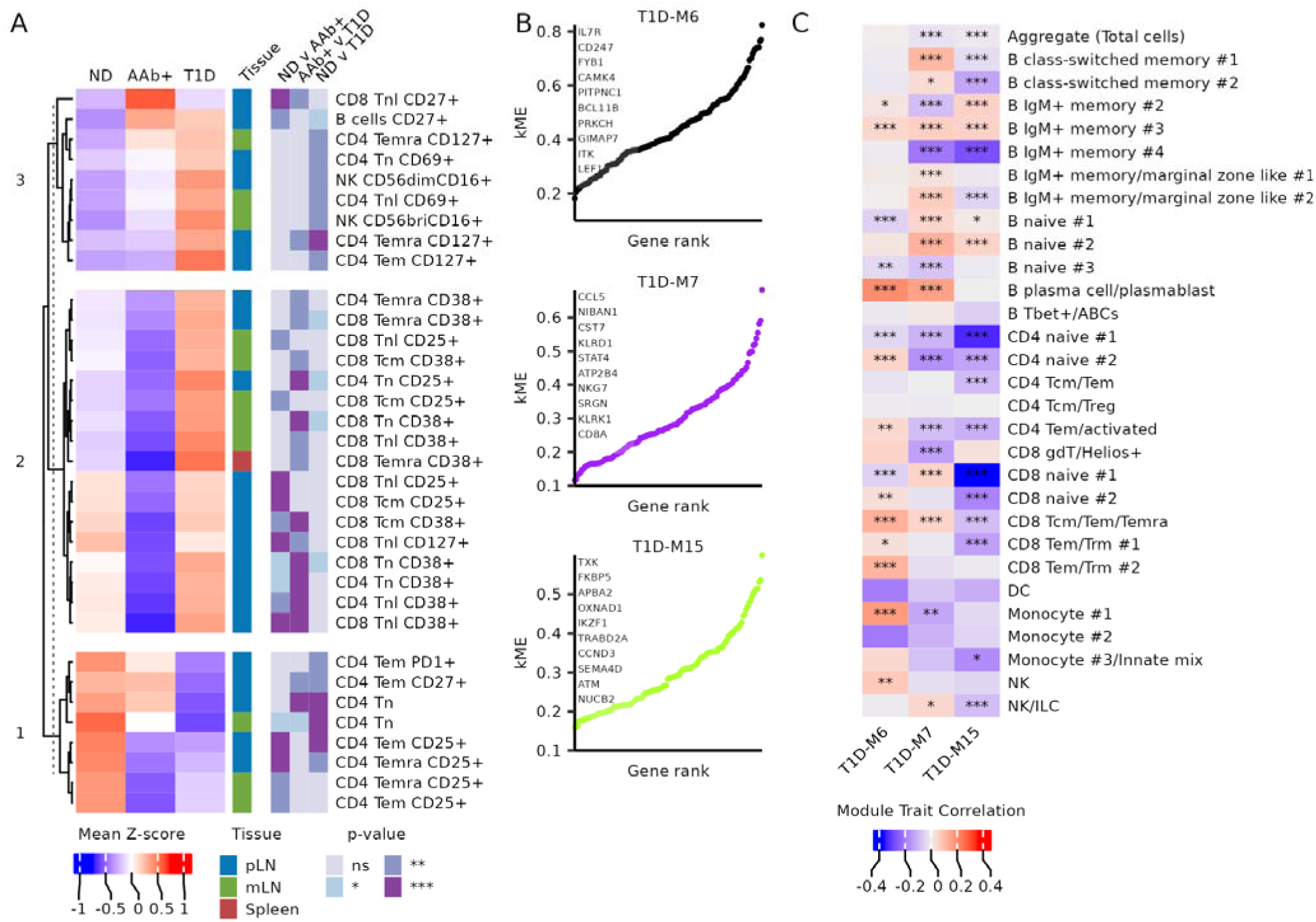
AAb+ and T1D associated shifts in immune phenotypes. (A) Heatmap of immune populations, detected by flow cytometry, from each tissue type that are significantly different in frequency in at least one comparison between disease states. Coloring of the heatmap represents the mean of Z-scores for a specific immune population within the specified tissue type. Statistical significance was calculated with robust one-way ANOVA, with post hoc testing using Hochberg’s multiple comparison adjustment. Only immune populations with a differential p-value < 0.01 in at least one disease comparison were plotted. (B) Modules of interest generated using WGCNA analysis on scRNAseq data from pLN lymphocytes in ND and T1D disease states. (C) Correlations of modules of interest between ND and T1D disease states, across all immune cell clusters. For all panels, * is p < 0.05, ** is p < 0.01, *** is p < 0.001.

We aimed to corroborate these flow cytometry findings using an unbiased weighted gene co-expression analysis (WGCNA) approach with the transcriptome modality of the CITEseq data to uncover covarying gene modules across T1D development. We only considered ND and T1D pLN in order to identify co-varying gene modules before and after overt autoimmunity. Seventeen gene modules covaried across the disease states, representing a diverse range of defined biological processes (Figure 2B-C, Extended Data Fig. 6A-B). Since most modules had weak correlations with T1D status, we focused on modules containing genes that overlapped with phenotypes observed via flow cytometry. Module 15 was significantly decreased in T1D within CD4+ and CD8+ naive T cell clusters (Figure 2B-C), similar to the decreased frequency of CD4+ Tn cells observed by flow cytometry (Figure 2A). Module 15 was anchored by *TXK*, a gene critical for TCR signaling^38^ and a regulator of interferon signaling^39^, and also contained the key lymphocyte development regulator *IKZF1* (Figure 2B). Module 7 included genes associated with proinflammatory signaling and cytotoxic immune cells including *CCL5*, *CST7^40^*, *KLRD1*, *STAT4^41^*, *NKG7*, and *SRGN* (Figure 2B). Module 7 was increased in the CD8+ Tcm/Tem/Temra and NK/ILC clusters (Figure 2C), similar to increased frequencies of cytotoxic CD56_dim_CD16+ NK cells seen in flow cytometry. Module 6, anchored by CD127 (*IL7R*), consisted of genes with known roles in T cell function and signaling such as *CD247*, *FYB1^42^*, *CAMK4^43^*, *PRKCH^44^*, *ITK,* and *LEF1* (Figure 2B). This module was positively correlated with T1D in CD4 Tem/activated clusters in the CITEseq dataset (Figure 2C), similar to the observed increase in frequency of CD127+ CD4+ Tem/TEMRA T cells in the flow cytometry dataset (Figure 2A). Overall, unbiased analysis of the CITEseq data generated from ND and T1D donor pLN lymphocytes can recapitulate some observations within the flow cytometry dataset. Subsequent analyses focused on significant phenotypes observed by flow cytometry that could be corroborated in the smaller CITEseq cohort.

### Loss and dysfunction of CD4+ Tregs in the pLN

Immune alterations before T1D onset are of particular interest to inform the development of early immune interventions capable of retaining β cell mass. In this regard, we observed a decreased frequency of CD25+ CD4+ Tem/Temra in the pLN of AAb+ donors that was maintained in the pLN of T1D donors (Figure 3A). CD4+ Tregs, phenotypically defined here by high CD25 and low CD127 surface expression, have previously been shown to be reduced in the pLN of T1D donors^35^, and certain single nucleotide polymorphisms (SNPs) in or nearby the CD25-encoding gene *IL2RA* are significantly associated with T1D risk^6,45^. Indeed, the frequency of CD4+CD25+CD127low Tregs was significantly reduced in both AAb+ and T1D pLN (Figure 3B-C), suggesting that CD4+ Tregs may be altered in the pLN prior to and during T1D onset in a tissue-restricted manner, as previously reported in the NOD mouse^46^. Using CITEseq, we further explored alterations in pLN Tregs within the CD4 Tcm/Treg cluster using a set of genes associated with various Treg signatures^47,48^. Core Treg genes *FOXP3*, *IKZF4*, and *IL2RA* were significantly reduced in the CD4 Treg/Tcm cluster in both AAb+ and T1D pLN (Figure 3D), showing that this signature appears prior to T1D onset. Other genes associated with Tregs, such as *CTLA4*, were increased in AAb+ versus ND pLN while genes negatively associated with Tregs such as *IL7R* decreased in AAb+ pLN, perhaps reflecting recent activation or attempted reconstitution of Tregs in AAb+ pLN. To discern changes within Tregs specifically, we analyzed transcriptional differences between disease states within *FOXP3*+ cells in the CD4 Treg/Tcm cluster (Figure 3E). Compared to ND Tregs, AAb+ Tregs expressed lower levels of *ANTXR2*, a gene critical for extracellular matrix interactions^49^, *GPR183*, a GPCR required for T cell positioning near activated T cells at the T cell zone-follicle interface^50^, and several *MAML* genes associated with Notch signaling^51^. Some of these genes also had decreased expression in T1D Tregs versus ND Tregs (Figure 3E). Compared to ND pLN Tregs, Tregs in T1D pLN expressed less *FOXP1*, a gene essential for Treg stability^52^, and *RUNX1*, which physically interacts with FOXP3 to upregulate Treg-associated genes^53^. To determine whether these signatures were unique to pLN Tregs, we examined the same signatures in mLN and splenic Tregs (Extended Data Fig. 5). We observed similar decreases in CD25+ CD4+ Tem/Temra frequency in AAb+ donors, but not T1D (Extended Data Fig. 5A-B). CD25+CD127low mLN CD4+ Tregs similarly decreased in mLN of AAb+ but not in T1D donors nor in spleen of any disease state (Extended Data Fig. 5C-D). Furthermore, we did not observe transcriptomic shifts in *FOXP3* or *IL2RA* seen in the pLN within mLN or spleen (Extended Data Fig. 5E). Together, these data indicate that pLN-, and to a lesser degree mLN- or splenic-, specific effects on Tregs occur both in frequency and gene expression associated with Treg lineage stability and function before T1D onset.

**Figure 3.**
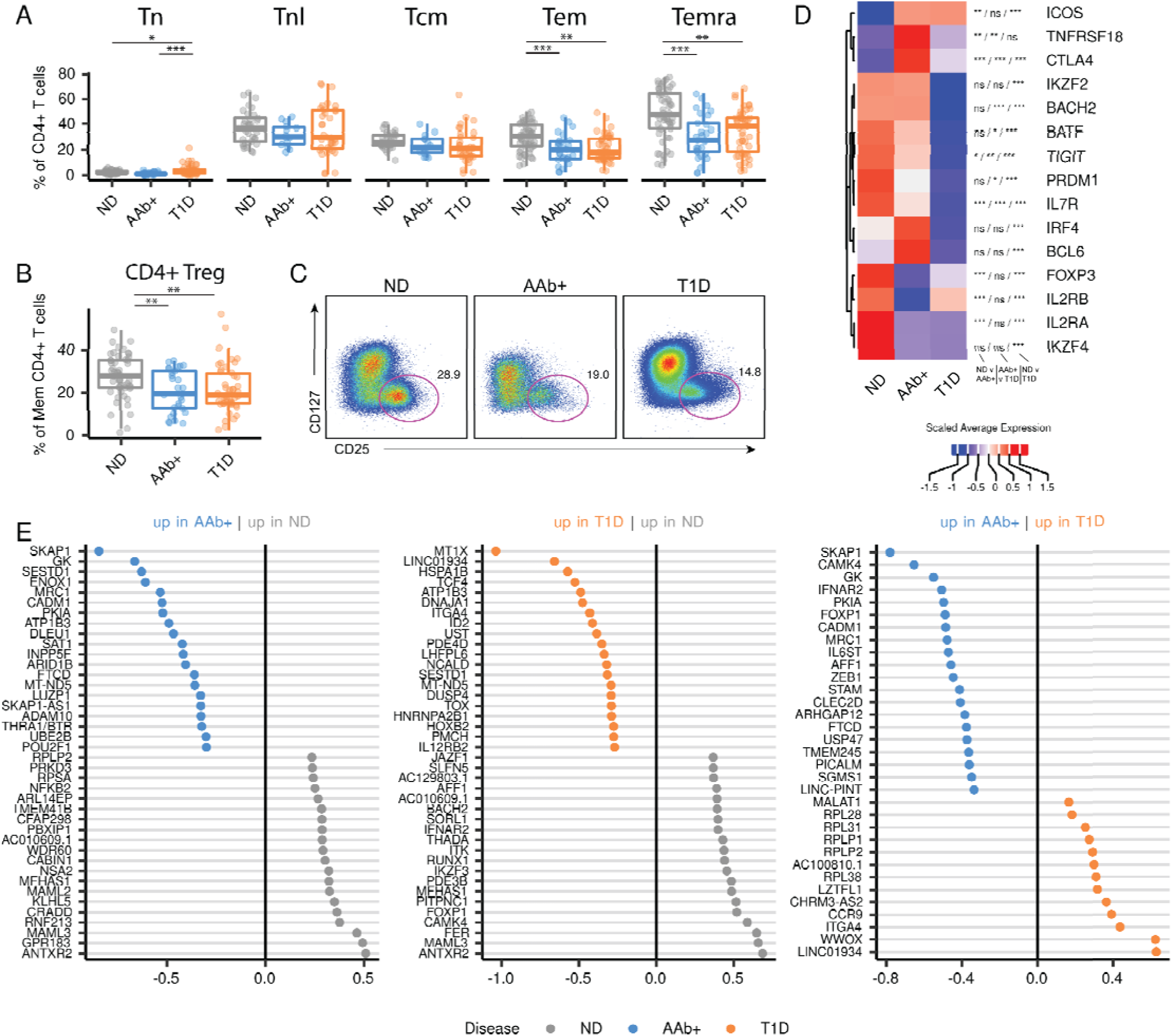
Loss and dysfunction of CD4+ Tregs in the pLN. (A) Frequency of pLN CD25+ cells or (B) CD25+ CD127- CD4+ Treg cells within CD4+ T cell subsets, as determined by flow cytometry. Statistical significance determined by robust ANOVA with post hoc testing using Hochberg’s multiple comparison adjustment. Boxplot represents median and interquartile range. (C) Representative two-parameter density plots of CD4+ Treg cells within the pLN, as measured by flow cytometry. (D) Expression of CD4+ Treg-associated genes within the CD4+ Treg/Tcm cluster from pLN. Expression scaled within each gene. Statistical significance determined by Wilcoxon Rank Sum Test and p-value adjustment using the Bonferroni method. (E) Top 20 differentially expressed genes (adjusted p value < 0.05, excluding common genes (see STAR Methods)) between disease states in *FOXP3*+ cells within the CD4 Treg/Tcm cluster in the pLN only. Statistical significance determined by Wilcoxon Rank Sum with genes with a log fold change threshold > 0.1 and p-value adjustment with the Bonferroni method across all combinatorial comparisons. For all panels in this figure, * is p < 0.05, ** is p < 0.01, *** is p < 0.001.

### Decreased naive T cell signatures in T1D pLN

We next characterized alterations in the naive CD4+ T cell pool in T1D donors. Naive CD4+ T cell frequency decreased in T1D versus ND pLN (Figure 4A) and mLNs (Extended Data Fig. 5G), while naive CD8+ T cell frequency in the pLN did not change with disease status (Figure 4B). Additionally, WGCNA module 15, which contained genes directly involved in T cell signaling and development, was negatively correlated with T1D status in both naive CD4+ and CD8+ T cells in T1D (Figure 2B-C, 4C). Donor age could not explain shifts in naive T cell phenotype, as age did significantly differ between disease groups, and the T1D group was slightly younger in the CITEseq cohort (Extended Data Table 1-2). Within the CITEseq data we identified two distinct clusters of naive CD4+ and CD8+ T cells: the first was highly enriched for naive T cell markers, termed naive #1, while the second had comparatively lower levels of naive surface markers and appeared activated, termed naive #2 (Extended Data Fig. 6). Both CD4 and CD8 naive #1 clusters in the pLN displayed a stronger negative correlation of WGCNA module 15 with T1D status, compared to their respective naive #2 cluster (Figure 4C). As a percentage of total pLN CD4+ T cells, CD4 cluster naive #1 trended lower in T1D compared to ND (Figure 4D), similar to that observed by flow cytometry (Figure 4A). Conversely, cluster CD4 naive #2 trended toward an increasing proportion of cells in T1D (Figure 4D). Neither of the naive CD8+ T cell clusters changed in frequency between disease states (Figure 4B, 4E). Several genes within WGCNA module 15 demonstrated significantly reduced expression in AAb+ or T1D pLN compared to ND pLN, particularly within the CD4 and CD8 Naive #1 clusters (Figure 4F-H). *TXK*, a gene critical for TCR signaling^38^ and a regulator of interferon signaling^39^, and *FKBP5*, a gene that coordinates with *FOXO4* to modulate cytokine production^54^, were significantly decreased in T1D versus ND across all naive subsets. *ATM*, a gene critical for DNA repair during V(D)J recombination^55^, was also decreased in both CD4 and CD8 naive #1 clusters but not naive #2 clusters from T1D donors. Taken together these data demonstrate a reduction in CD4+ naive T cells in T1D that is accompanied by decreased expression of genes in both CD8+ and CD4+ naive T cells involved in processes essential for T cell signaling and DNA repair.

**Figure 4.**
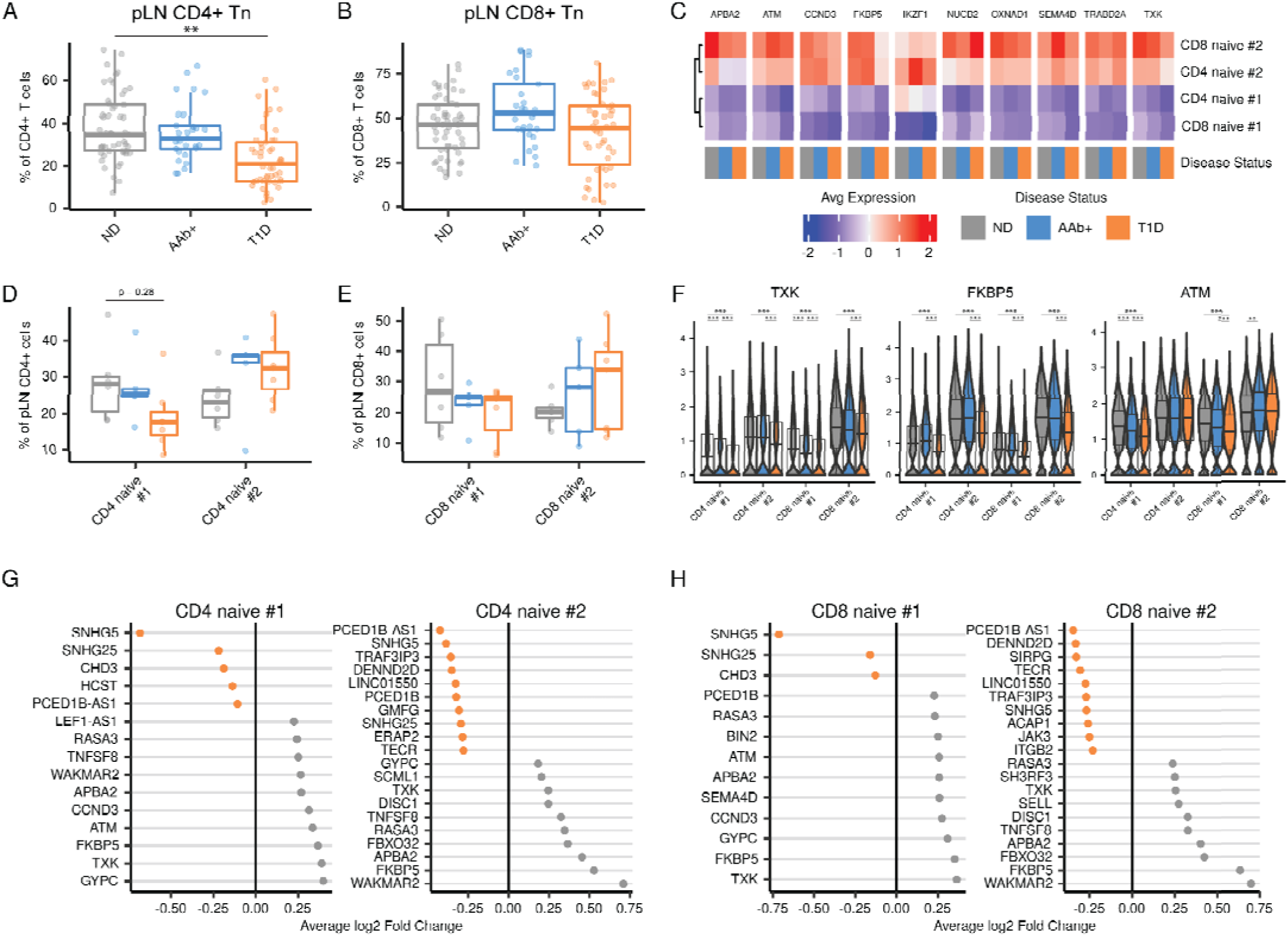
Decreased naive T cell signatures in T1D pLN. A) Frequency of pLN CD4+ Tn within total CD4+ T cells or (B) pLN CD8+ Tn within total CD8+ T cells, as determined by flow cytometry. Statistical significance determined by robust ANOVA with post hoc testing using Hochberg’s multiple comparison adjustment. Boxplot represents median and interquartile range. (C) Mean normalized expression of the top 10 most inter-connected genes in WGCNA module 15, plotted across the naive T cell clusters in the pLN and disease state. (D) Frequency of the two naive CD4+ T cell clusters and (E) the two naive CD8+ T cell clusters in the pLN, across disease states. Boxplot represents median and interquartile range. P value determined by Wilcoxon ranked sum test with Benjamini-Hochberg multiple test correction. (F) Normalized expression of genes of interest from WGCNA module 15, from the pLN only. Statistical significance determined by Wilcoxon Rank Sum Test with a log fold change threshold of 0.1 and p-value adjustment using the Bonferroni method. (G) Differentially expressed WGCNA module 15 genes CD4 Naive #1 and CD4 Naive #2 clusters and (H) CD8 Naive #1 and CD8 Naive #2 clusters from the pLN, between ND and T1D donors. Statistical significance tested using Wilcoxon Rank Sum Test with p-value adjustment using the Bonferroni method. For all panels in this figure, * is p < 0.05, ** is p < 0.01, *** is p < 0.001.

### Memory CD8+ T cells display a stem-like phenotype in AAb+ and T1D pLN

As CD8+ T cells are widely thought to play a direct role in human T1D pathology by infiltrating islets and eliminating β cells^1,28^, we sought a deeper examination of CD8+ T cell perturbations before and after T1D onset. In human pLN, we found multiple CD8+ T cell phenotypic changes before and after T1D onset in both flow cytometry and CITEseq datasets. We observed a marked decrease in the proportion of cells expressing the activation markers CD25 (Figure 5A) and CD38 (Figure 5B) in AAb+ donors compared to ND. Decreased CD25 expression was noted amongst all memory CD8+ T cell subsets in AAb+ donor pLNs, and otherwise only amongst CCR7- CD8+ memory T cell populations in mLN (Extended Data Fig. 5I-J). The CD38+ frequency decreased in CD8+ Tn, Tn-like (Tnl), and Tcm subsets and trended lower in CD8+ Tem and Temra subsets in the pLN of AAb+ donors. In mLN and spleen, CD38 surface expression only decreased in splenic Tnl of AAb+ donors compared to ND. Overall, the frequency of CD25+ and CD38+ CD8+ T cell subsets decreased in the pLN, and to a lesser degree in the mLN and spleen, of AAb+ donors in comparison to ND.

**Figure 5.**
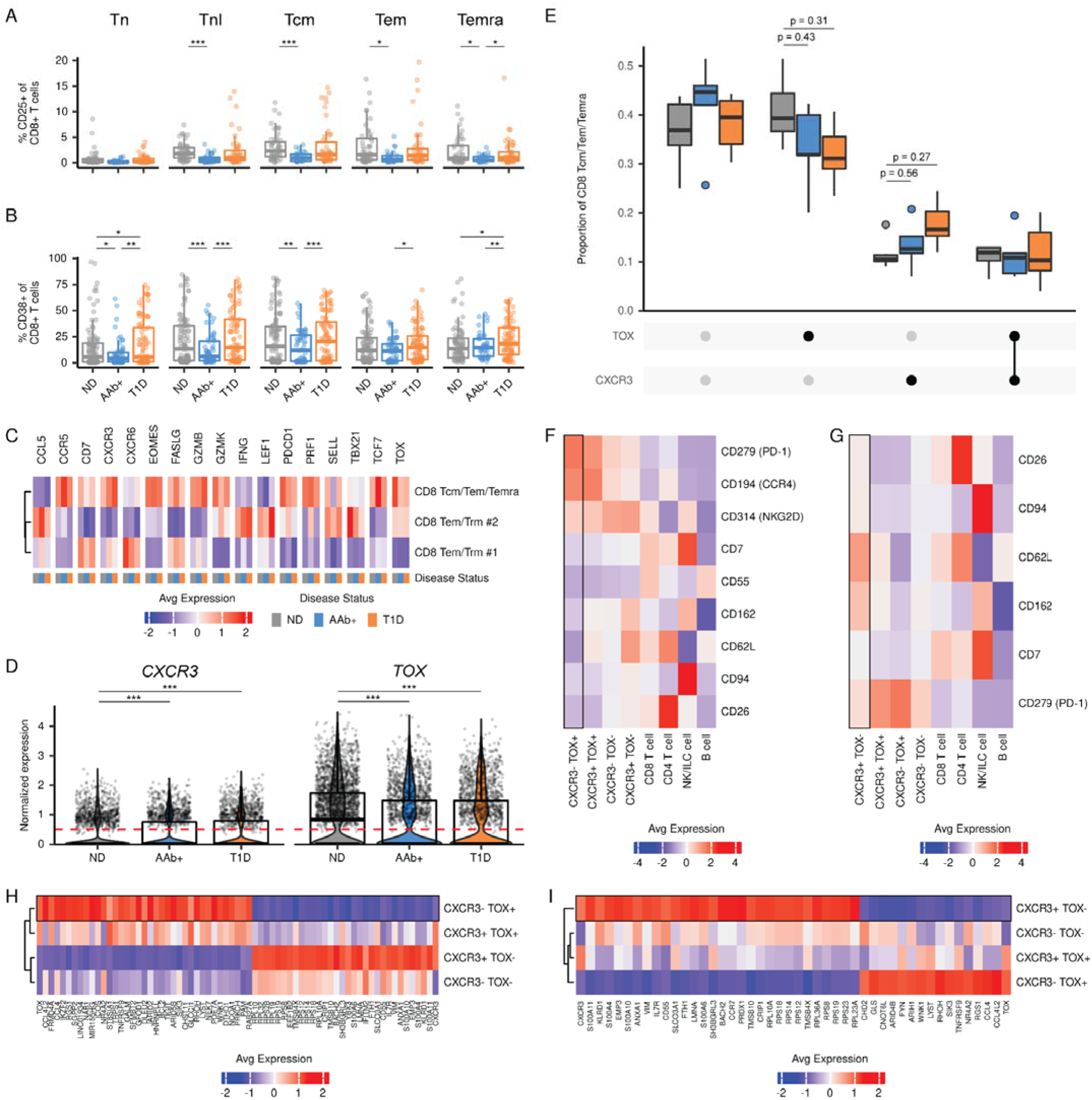
Memory CD8+ T cells display a stem-like phenotype in AAb+ and T1D pLN. (A) Frequency of pLN CD25+ cells or (B) CD38+ cells within pLN CD8+ T cell subsets, as determined by flow cytometry. Statistical significance determined by robust ANOVA with post hoc testing using Hochberg’s multiple comparison adjustment. Boxplot represents median and interquartile range. (C) WGCNA module 7 and effector CD8+ T cell genes of interest mean normalized expression in the pLN, across disease state. (D) Normalized expression of *CXCR3* and *TOX* in CD8 Tcm/Tem/Temra cells in the pLN. Statistical significance determined by Wilcoxon Rank Sum test with Bonferroni correction done between all combinatorial tests. Dashed red line indicates the exclusive threshold (> 0.5) for demarcating positive gene expression for later plots. (E) Frequency of pLN CD8 Tcm/Tem/Temra cells expressing permutations of *CXCR3* and *TOX*, across disease state. P value determined by Wilcoxon rank sum test with Benjamini-Hochberg multiple test correction. (F) Differential surface protein ADTs enriched in *CXCR3*-*TOX*+ cells and (G) *CXCR3*+*TOX*- cells compared to all other permutations of *CXCR3* and *TOX* expression within CD8 Tcm/Tem/Temra cluster in the pLN. Total CD8+ T cells, total CD8+ T cells, the NK/ILC cell cluster, and total B cells were used as comparators. Surface markers are ranked in descending order of fold change between *CXCR3*-*TOX*+ cells and *CXCR3*+*TOX*- cells. Statistical significance determined by Wilcoxon rank sum test on ADTs with log fold change threshold > 0.1 followed by Bonferroni multiple test adjustment (adjusted p value < 0.05 denotes significant differential expression). Populations of interest highlighted by the black outline. (H) Genes differentially expressed in *CXCR3*-*TOX*+ cells and (I) *CXCR3*+*TOX*- cells compared to all other permutations of *CXCR3* and *TOX* expression within CD8 Tcm/Tem/Temra cluster in the pLN. Genes are ranked in descending order of fold change between *CXCR3*-*TOX*+ cells and *CXCR3*+*TOX*- cells. Statistical significance determined by Wilcoxon rank sum test with log fold change threshold > 0.25 followed by Bonferroni multiple test adjustment (adjusted p value < 0.05). Only genes with a p < 0.0001 are plotted. Populations of interest highlighted by the black outline. For all panels in this figure, * is p < 0.05, ** is p < 0.01, *** is p < 0.001.

We further explored T1D-associated memory CD8+ T cell alterations in relation to disease state using CITEseq (Figure 5C). CD25 and CD38 ADT signatures could not be used to corroborate phenotypes observed with flow cytometry due to low resolution of these markers in memory CD8+ T cell clusters. Therefore, we instead utilized genes within the proinflammatory and cytotoxicity associated WGCNA Module 7 that positively correlated with T1D in memory CD8+ T cells (Figure 2B-C) and genes associated with autoimmune or tissue-targeting CD8+ effector memory T cells (Figure 5C)^34,56^. The cells in the CD8 Tcm/Tem/Temra cluster included a diverse range of memory CD8+ T cell subsets, based on gene expression signals associated with stem, effector, and effector memory CD8+ T cells such as *TCF7*, *EOMES*, *TOX*, *PRF1*, *GZMB*, and *CXCR3* (Figure 5C). Of these genes, *CXCR3* and *TOX* were the only genes differentially expressed in both AAb+ and T1D pLN relative to ND pLN, with expression of *CXCR3* increasing and *TOX* decreasing (Figure 5D). *TOX* had similar expression patterns in mLN and spleen, while *CXCR3* expression decreased in T1D versus ND mLN (Extended Data Fig. 5M-N), suggesting that the pLN may have a unique *CXCR3* gene expression pattern compared to the other tested tissues (Extended Data Fig. 5M-N). The frequency of *CXCR3*-*TOX*+ and *CXCR3*+*TOX*- cells in the pLN trended in the same patterns as *TOX* and *CXCR3* gene expression (Figure 5E), suggesting that the cells driving this gene expression pattern expressed either *CXCR3* or *TOX*. Compared to all cells in the CD8 Tcm/Tem/Temra cluster, *CXCR3*-*TOX*+ cells had higher levels of PD1 and CCR4 surface protein (Figure 5F). Conversely, *CXCR3*+*TOX-* cells expressed relatively higher levels of the lymph node homing receptor CD62L (L-selectin), the immune inhibitory receptor CD94 (*KLRD1*), and the costimulatory molecule CD26 on the cell surface (Figure 5G). Within the CD8 Tcm/Tem/Temra cluster, *CXCR3*-*TOX*+ cells and *CXCR3*+*TOX-* cells had distinct gene expression patterns (Figure 5H-I). Genes with higher expression in *CXCR3*-*TOX*+ cells included chemokines *CCL4* and *CCL4L2*, inhibitory receptor *TIGIT*, costimulatory molecule *ICOS*, and transcription factors *IKZF2* and *BCL2* (Figure 5H). *CXCR3*+*TOX-* cells expressed higher levels of immunosuppressive genes *KLRD1* and *CD55*, the CD127-encoding gene *IL7R*, and the stemness-enforcing transcription factor *BACH2*^57,58^. In total, pLN CD8+ T cells exhibited changes that manifested in AAb+ donors prior to T1D development: decreased cell surface expression of activation markers CD25 and CD38, a reduction of an effector *TOX*+ population, and an increase in a stem-like *CXCR3*+ population. The mLN and spleen also contained some, but not all, of these phenotypic differences between disease states, specifically overlapping with pLN in CD25 surface expression and *TOX* expression patterns.

### Cytotoxic NK cells more prominent in T1D pLN

Finally, we examined innate cell populations for potential changes associated with AAb+ or T1D status. While few overt differences were noted within most innate cell lineages present within the pLN or mLN across the different cohorts, we found that the cytotoxicity-associated WGCNA module 7 (Figure 2B-C) was positively correlated with T1D status in pLN N K cells (Figure 2A), and there was a proportional increase in the cytotoxic CD56_dim_CD16+ NK cell frequency in T1D pLN (Figure 6A-B). We further assessed differential gene expression within NK cell clusters between ND and T1D, focusing on genes within WGCNA module 7 and additional NK cell cytotoxicity associated genes from the KEGG pathway database^59–61^. *GZMB*, *CRTAM*, *IFNG*, and other genes associated with NK cell cytotoxicity were upregulated in T1D (Figure 6C-D). Further, the NK cell inhibitory receptor *KLRB1* was the most downregulated gene by fold change in the specific gene set (Figure 6C,E). *GZMB*+ NK cells in the pLN were highly enriched in genes associated with cytotoxicity such as *NKG7*, *KLRD1*, and *GNLY* (Figure 6F), while *GZMB*- NK cells in the pLN expressed *IL1R1* and the immunosuppressive enzyme *IL4I1^62^*(Figure 6G). Compared to ND, *GZMB* expression was not increased in T1D in the mLN or spleen, while *KLRB1* decreased in expression in the mLN (Extended Data Fig. 5O-P), indicating that the increased cytotoxic NK cell frequency and transcriptomic signature was observed in the pLN, and not the mLN or spleen, of T1D donors.

**Figure 6.**
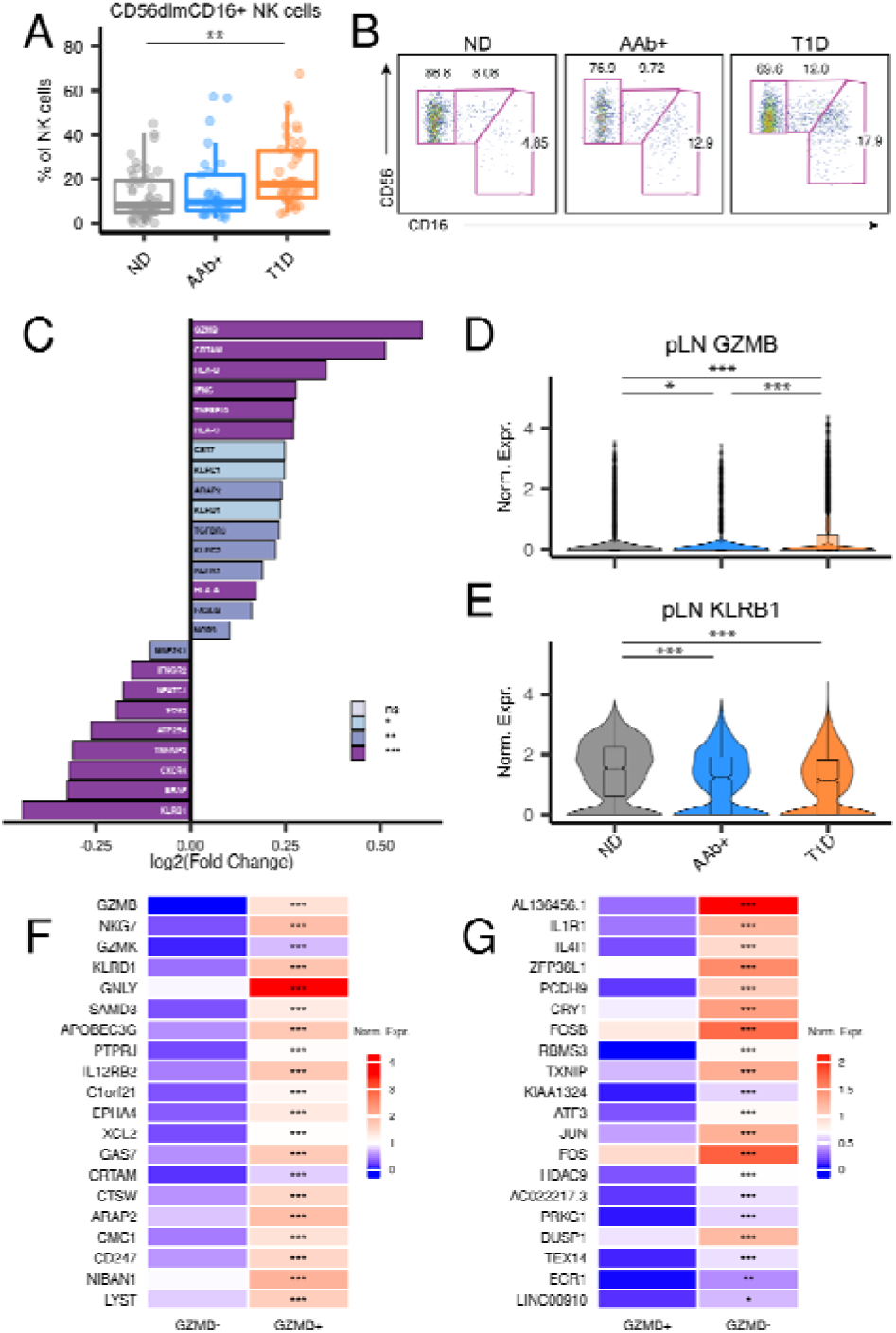
Cytotoxic NK cells more prominent in T1D pLN. (A) Frequency of CD56_dim_CD16+ NK cells within the pLN, as measured by flow cytometry. Statistical significance determined by robust ANOVA with post hoc testing using Hochberg’s multiple comparison adjustment. Boxplot represents median and interquartile range. (B) Representative two-parameter density plots images of NK cell subsets within the pLN. (C) List of differentially expressed genes, ranked by descending log2 fold change, of genes that are significantly more expressed in T1D pLN. Cells are from a combination of the NK and NK/ILC clusters, from the pLN only. Gene sets to test came from the Module 7 of the WGCNA analysis, which was significantly increased in NK cell clusters in T1D, and from the “Natural killer cell mediated cytotoxicity” gene set from Kyoto Encyclopedia of Genes and Genomes (KEGG). (D) Normalized expression of *GZMB* and (E) *KLRB1* within the combined NK and NK/ILC clusters in the pLN. Statistical significance tested using Wilcoxon Rank Sum Test with p-value adjustment using the Bonferroni method. (F) Mean normalized expression of genes significantly increased in *GZMB*+ and (G) *GZMB*- cells from the combined NK and NK/ILC clusters in the pLN. P value determined by Wilcoxon Rank Sum Test with p-value adjustment using the Bonferroni method. For all panels in this figure, * is p < 0.05, ** is p < 0.01, *** is p < 0.001.

### Immune populations correlate with HLA genetic risk

HPAP collects and publicly shares extensive clinical information on each donor, allowing for the correlation of immune phenotypes with these metadata. One such parameter is HLA type, where certain alleles of HLA-DR and HLA-DQ convey significant risk for T1D^63–65^. Higher genetic risk is associated with rapid onset of T1D^66^ and can distinguish between young adults with T1D versus T2D^67^. Therefore, profiling the immune system in individuals with high genetic risk may uncover immune perturbations associated with T1D disease susceptibility. Using only the HLA risk score portion of Genetic Risk Score 2 (GRS2)^68^, we generated an HLA genetic risk score (HLA-GRS) for each donor that accounts for HLA DR-DQ haplotype interactions^17,68^, with a higher score indicating a higher risk for disease development. As expected, T1D donors had increased HLA-GRS compared to ND (Figure 7A). After controlling for the effects of disease state on immune population frequency, we observed moderate correlations between HLA-GRS and the frequency of specific pLN immune populations. For example, when grouping ND and AAb+ donors, the frequency of CD8+ Tcm and Tem expressing CXCR5, HLA-DR and/or CD38 were positively correlated with HLA-GRS (Figure 7B-D). When grouping ND and T1D donors, similar populations were associated with HLA-GRS but did not reach statistical significance (data not shown), likely due to these populations being regressed out due to their increased frequency in T1D pLNs (Figure 2A). When binning AAb+ and T1D donors, the frequency of CD69+ cells in B cells and memory T cell subsets correlated positively with HLA-GRS (Figure 7E-F). The frequency of CD127+ memory T cell subsets also positively correlated with HLA-GRS, with a reciprocal decrease in the frequency of CD127-T cells (Figure 7E,G). Together, these data indicate that higher risk HLA haplotype combinations are positively associated with the frequency of certain pLN immune populations, particularly activated CD8+ memory T cells, CD127+ T cells, and CD69+ B and T cells.

**Figure 7.**
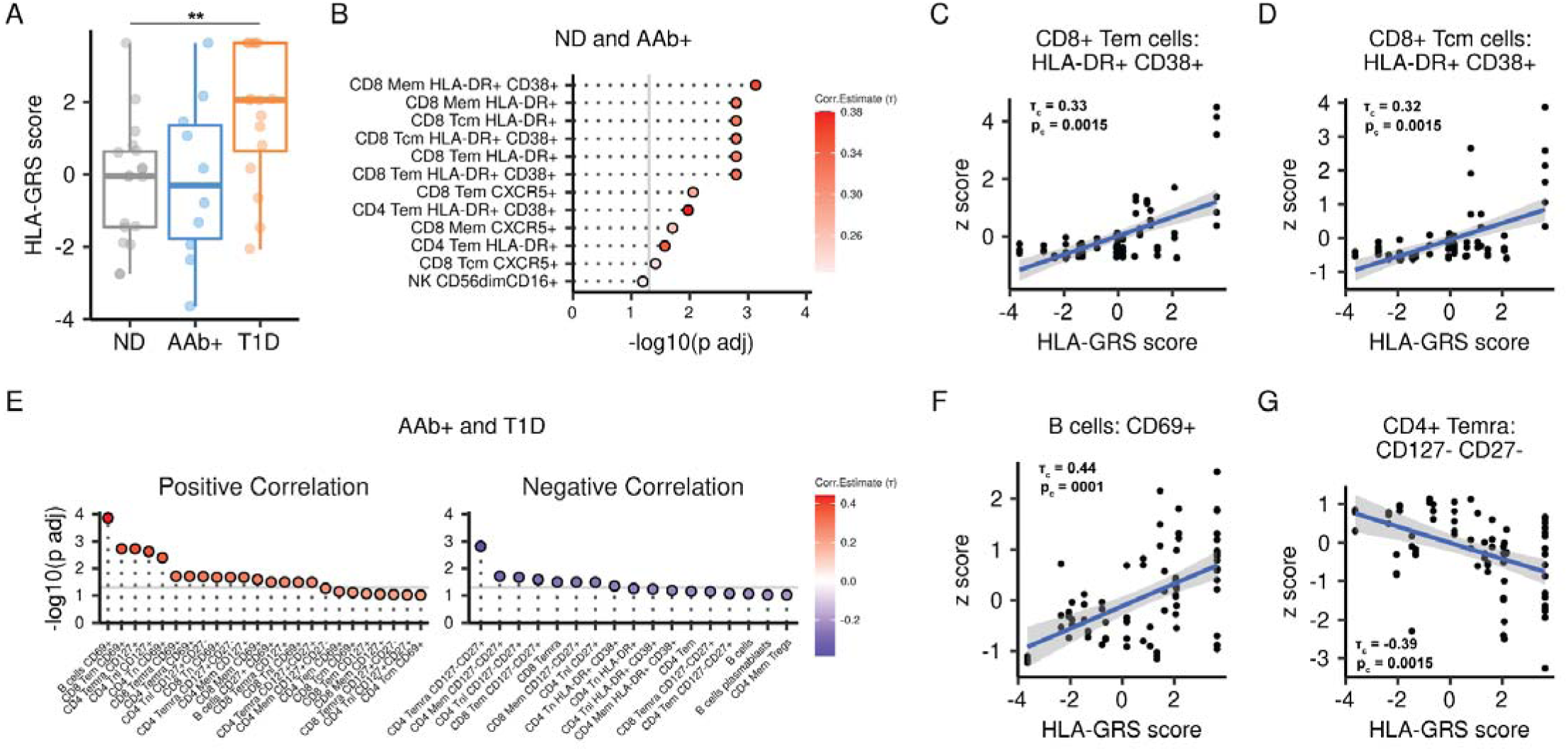
Immune populations correlate with HLA genetic risk. (A) HLA-GRS score of donors in the cohort. Boxplot represents median and interquartile range. P value generated with Dunn’s test with multiple hypothesis correction adjustment using Holm’s method. * is p < 0.05. (B) Immune populations that significantly correlate with HLA-GRS in ND and AAb+ donors. Grey line represents an adjusted p value ≥ 0.05. Dot fill represents a Kendall tau correlation value corrected for disease state effects. (C) and (D) Representative plots of HLA-GRS versus immune population frequency in ND and AAb+ donors. τ_c_ is the Kendall tau correlation value corrected for disease state effects, represented by the blue linear regression line with standard error in grey. p_c_ is the p value adjusted for disease state effects and corrected with Benjamini-Hochberg multiplicity adjustment. (E) Immune populations that significantly correlate with HLA-GRS in AAb+ and T1D donors. All plot parameters follow (B). (F) and (G) Representative plots of HLA-GRS versus immune population frequency in AAb+ and T1D donors. All plot parameters follow (C) and (D).

## Discussion

Our current knowledge of the landscape of immune perturbations during the ontogeny of T1D largely originates from the NOD mouse model and targeted analysis of specific immune subsets in human peripheral blood, with limited studies of human pLN^32,33,35^. Here, we broadly profiled lymphocyte subsets from lymphatic tissues in a large cross-sectional cohort of ND, AAb+, and T1D human organ donors to define immunological perturbations occurring in pLN prior to and after disease diagnosis. Compared to ND, AAb+ and T1D pLN have a reduced Treg signature and increased stem-like *CXCR3*+ CD8+ T cells. Some perturbations were T1D pLN specific, including a reduced naive T cell signature and an increased frequency of cytotoxic CD56_dim_CD16+ NK cells. Additionally, several immune populations in the pLN correlated with HLA-GRS independently of the effects of disease status, in particular CD8+ T cell activation signatures and increased CD69 and CD127 surface expression on T cells and/or B cells. Importantly, immune alterations were most readily observed in lymphoid tissues that drain the pancreas as opposed to the spleen, implying that disease-related effects may be diluted or influenced by other factors at sites more distal from autoimmune inflammation.

One of the clearest observed signatures was a marked decrease in CD25 expression on CD4+ and CD8+ T cells in the pLN of AAb+ individuals. This was most strongly reflected by an overall decrease in CD4+ Tregs, a phenomenon observed previously in T1D^35^, but shown here to also manifest in AAb+ donors and in individuals with T1D. IL-2 signaling and production deficiencies have long been tied to T1D susceptibility, as PBMCs from children with recent onset T1D produce less IL-2^69,70^, and several T1D susceptibility genomic loci are involved in IL-2 signaling^6,45^. IL-2 induced signaling responses are critical to maintaining T cell tolerance to self-antigen through the maintenance of CD4+ Tregs^71^, and IL-2 signaling strongly influences CD8+ memory T cell differentiation^72^. Decreases in pLN Treg frequency in AAb+ donors highlight the evolution towards an inflammatory environment that may occur in pLN before T1D onset. Further, while CD25+ CD4+ Tem frequency in the mLN and spleen have similar patterns across disease states compared to pLN, a decline of a Treg signature in AAb+ or T1D disease states was not observed in mLN or spleen, indicating that the pLN are uniquely losing this cell population known to be critical for autoimmune control.

The loss of Tregs in AAb+ individuals was accompanied by an increase of a stem-like CD8+ T cell population closely resembling one found in multiple autoimmune contexts, including a population that resides in the pLNs and drives autoimmunity in NOD mice^34^ and is associated with ulcerative colitis in mice and humans^73^. In humans, stem-like CD8+ T cells^12^ and CXCR3+ CD8+ T cell subsets^17^ in PBMCs are positively associated with T1D status. In human AAb+ and T1D pLN, we also observed increased frequencies of less differentiated stem-like *CXCR3*+ memory CD8+ T cells. Importantly, *CXCR3*+ memory CD8+ cells did not change across disease state in the mLN, suggesting localization of this effect to the pLN. We further found that a *TOX*+ memory CD8+ T cell population defined by surface PD1 and CCR4 protein and gene expression of *CCL4*, *IKZF2,* and *BCL2* reciprocally decreased in the pLN of AAb+ individuals. While PD1+ CD8+ T cells expressing *TOX* have been described as being functionally exhausted in some contexts, many polyfunctional non-exhausted CD8+ T cells express PD1 and TOX in human lymph nodes^74^. The PD1+ *TOX*+ CD8+ T cell population shares features with *IKZF2+*KIR+ CD8+ T cells, which play a role in peripheral tolerance during inflammatory conditions^75^. It is unclear how these events are temporally associated with progression as T1D-associated AAb specificities develop^76^, but a loss of Treg activity or changes in IL2 signaling could enable the survival and expansion of T1D autoantigen-specific CD8+ T cells in the local pLN environment.

We also observed an increased proportion of cytotoxic NK cells, and decreases in frequency and transcriptomic alterations in naive CD4+ and CD8+ T cells, in pLN and mLN of T1D donors. Although various pathways may explain the general differentiation of naive T cells and NK cells, evidence in the context of T1D implicates cytokine signaling, and in particular γ-chain cytokines, as a potential mechanism behind these observations. Long-term inflammation drives immune aging, activation, and differentiation signatures in PBMCs of T1D donors^17^. γ-chain cytokines IL7 and IL21 are elevated in the circulation of long-standing, and not newly diagnosed, T1D donors^77,78^, and IL7 and IL21 signaling are required for diabetes development in NOD mice^79–81^. IL21 and IL15 promote NK cell differentiation into cytotoxic subsets including CD56_dim_CD16+ NK cells^82^, and γ chain cytokines promote naive T cell differentiation^83^, similar to phenotypic changes we observed in these immune subsets. Additionally, IL7 is known to increase the expression of its canonical receptor CD127^84^, and we observed CD127 expression in the pLN was positively associated with T1D status and HLA-associated T1D risk. Several IL2-related phenotypic changes were identified in the pLN of AAb+ donors, implying that certain γ chain cytokines may impact earlier stages of T1D progression and others after disease establishment. Examining if γ chain cytokine signals actively drive the observed phenotypes in the T1D context, and whether they derive from the lymph node or upstream from the pancreas, will inform therapeutic strategies for immune intervention during or after T1D onset.

By calculating T1D genetic risk conveyed by HLA class II alleles, we found that the frequency of memory CD8+ T cells with HLA-DR and CD38 surface expression was positively correlated with HLA-GRS, indicating that non-T1D individuals with higher-risk HLA alleles have an increased frequency of activated memory CD8+ T cells in the pLN. While cytotoxic CD8+ T cells infiltrate islets and eliminate β cells in T1D^26,29^, it is unclear how or where these autoimmune CD8+ T cells develop in humans. Evidence from the NOD mouse implicates stem-like autoimmune CD8+ T cells as a reservoir of T cells that infiltrate the pancreas^34^, and we observed an expansion of stem-like memory CD8+ T cells in the pLN of AAb+ and T1D donors. It is unknown whether pLN CD8+ T cell activation observed in high-risk individuals is related to the expansion of stem-like CD8+ T cells, but a higher basal T cell activation state may potentiate expansion of autoimmune CD8+ T cells. We found that a higher HLA-GRS in AAb+ and T1D donors positively correlates with CD127 surface expression on T cells. CD127, along with the common γ chain receptor subunit, comprises the IL7R that is critical for T cell homeostatic proliferation^85^, implying that pLN T cells in individuals with higher HLA-GRS may have heightened IL7 responsiveness^86^. AAb+ and T1D individuals with a higher HLA-GRS also had increased frequencies of CD69+ B and T cell subsets, indicating these individuals may have more tissue resident^87^ or activated^88^ immune cells. As tissue residency is critical for local tissue immune memory responses^87^, further investigation is needed to explore if high risk individuals have an expanded immune memory pool that could potentiate autoimmune responses.

Our study was limited in a number of aspects. First, due to the complexities of donor acquisition, we were limited in our ability to acquire enough AAb+ organ donors with two or more T1D-associated AAbs, preventing a properly powered subanalysis of single versus multiple AAb+ donors. As the study design is inherently cross-sectional, we do not have longitudinal data, including glucose tolerance tests, to properly stage AAb+ individuals. This limits our ability to stage the progression of immune perturbations that we observed in our cross-sectional study. Individuals without diabetes who have multiple T1D-associated AAbs are rare, and generally progress to T1D more rapidly than those with single autoantibodies^2^. Furthermore, as not all single AAb+ donors develop T1D^2,89^, additional studies of pLN from donors with >1AAb+ across a range of AAb specificities will be necessary to define immunological associations with pre-T1D. Second, due to blood samples being drawn well before donor organ harvest, we determined that peripheral blood lymphocytes yielded unreliable flow cytometric and transcriptomic data (data not shown). As such, we cannot directly compare our results to previous studies focusing exclusively on peripheral blood. Third, we restricted our flow cytometric analysis to freshly isolated lymphocytes, rather than frozen lymphocytes, due to evidence of cryopreservation-induced changes in certain cell surface markers^90^. This limits subsequent flow based analysis to prospectively acquired cohorts. Fourth, we were unable to simultaneously assess B cell or T cell repertoires in the CITEseq analysis due to technical limitations from 3’ mRNA transcriptomic analysis. Hence, studies of T cell or B cell T1D antigen specificity are pending.

The deep immune profiling of rare and valuable tissue samples from organ donors collected across the spectrum of disease stages reported here provide an expansive resource for both the T1D community as well as individuals interested in lymph node immunology. This resource is structured as an immune cell atlas in an open data format with the goal of facilitating novel investigative approaches to understand disease mechanisms and identify novel targets to intervene in T1D development. Importantly, the conjunction of the high-dimensional analyses allowed for the observation of subtle immune changes associated with AAb positivity and T1D status. The datasets provide foundational immunobiology information for the research community that could help uncover potential therapeutic targets to delay, prevent, or ameliorate T1D.

## Acknowledgements

Funding: UC4-DK-112217 (to K.H.K., T.M.B., E.T.L.P., A.N., and M.R.B.), Penn IDOM Internal Pilot Grant 5-P30-DK-019525 (to M.R.B.), JDRF 3-SRA-2022-1237-S-B (to M.R.B.), JDRF PDF 3-PDF-2023-1323-A-N (to G.J.G.), Human Islet Research Network (HIRN). This manuscript used data acquired from the Human Pancreas Analysis Program (HPAP-RRID:SCR_016202), and the Human Islet Research Network (RRID:SCR_014393) consortium (UC4-DK-112217, U01-DK-123594, UC4-DK-112232, and U01-DK-123716). This research was performed with the support of the Network for Pancreatic Organ donors with Diabetes (nPOD; RRID: SCR_014641), a collaborative type 1 diabetes research project sponsored by the Juvenile Diabetes Research Foundation (JDRF) (nPOD: 5-SRA-2018-557-Q-R), and The Leona M. & Harry B. Helmsley Charitable Trust (grant 2018PG-T1D053). Genetic risk scoring data was analyzed with support from NIAID P01 to T.M.B. (AI042288). Flow cytometry data was generated in the Penn Cytomics and Cell Sorting Shared Resource Laboratory at the University of Pennsylvania (RRID: SCR_022376), which is partially supported by the Abramson Cancer Center NCI Grant (P30 016520).

Personnel: We thank K. Trihemasava, J. Shoush, and J.M.L Nordin for their assistance processing and running flow cytometry on HPAP donors, the Betts lab for their support with annotating the CITEseq dataset, and B.M.W. Golden, B.B. Golden, and B. Golden for their support.

## Contributions

G.J.G., A.S.J, M.B.P., L.K.C., and M.R.B designed flow cytometry experiments. G.J.G., A.S.J, M.B.P., L.K.C., and J.T.H. performed flow cytometry experiments. G.J.G. analyzed the flow cytometry data. G.J.G., V. W., J.T.H., K.A., A.S.J., C.L., and J.S.G. processed HPAP tissue samples. G.J.G., V.W., and M.R.B. designed the CITEseq experiments. G.J.G performed the CITEseq experiments. G.J.G, V.W., J.T.H, M.B.P., J.J.K., E.T.L.P., and M.R.B. annotated the CITEseq dataset. G.J.G, V.W., K.A., and M.R.B. analyzed and interpreted the CITEseq dataset. M.R.S. generated the HLA-GRS. J.T.H. created tables summarizing donor information. G.J.G., V.W., and K.A. performed statistical analyses. M.A.A., K.H.K., and A.N. oversaw subject selection, clinical data acquisition, clinical data analysis, research data reporting for study subjects, and editing the manuscript. G.J.G., V.W., T.M.B, E.T.L.P., A.N., and M.R.B formulated the ideas and direction of the project. G.J.G., V.W., and M.R.B wrote the manuscript.

## Ethics Declarations

The authors declare no competing interests.

## Methods

### Tissue collection

Experimental model and study participant methods are described^29,91^. Details on the samples obtained through the HPAP are available online at https://hpap.pmacs.upenn.edu and described^37^. Whole organ samples are obtained via the local organ procurement organization or via University of Florida/nPOD. The consent for procurement of research pancreata was obtained by staff of the organ procurement organization. All human samples were de-identified and IRB exempt. Donors were evaluated for autoantibody status (GAD, IA-2, IAA and ZnT8), C-peptide levels, and underwent clinical chart review. Clinical data including diagnosis of T1D, HbA1c, BMI, age, gender, HLA type, and more are recorded in the HPAP database. Additional data from histology, immunophenotyping, repertoire profiling, metabolic and transcriptomic studies are generated on all HPAP donors, providing a shared and publicly available comprehensive resource.

### Tissues and cells

To isolate pancreatic and mesenteric LN lymphocytes, LNs obtained from brain-dead organ donors were submerged in 5 mL R10 media (RPMI + 10% FBS + 1% penicillin/streptomycin + 2mM L-glutamine) supplemented with 10U/mL DNase I (Roche), cleaned of visceral fat, and cut into small pieces (∼2mm x ∼2mm). Tissue fragments were placed into a sterile 70 μm mesh (Miltenyi Biotech) and pushed through the mesh with the blunt end of a sterile 5 mL syringe plunger. Residual tissues and cells in the mesh were washed with 45 mL of R10 + DNAse, centrifuged at 500 xg for 7 minutes, resuspended in 10 mL of R10 + DNAse, and counted. To isolate splenocytes, approximately 30 g of spleen tissue was placed in R10 supplemented with 10U/mL DNase I and 3 mg/ml collagenase D (Sigma), the capsule was removed, and remaining tissue was cut into 4mm x 4mm pieces. The tissue was then mechanically dissociated using a GentleMACS (Miltenyi Biotech) and incubated at 37 C for 15 min with gentle inversion. After digestion, the cell suspension was filtered through a 100 μm strainer, and erythrocytes were lysed by ACK buffer (Corning). Post-ACK, cells were resuspended in R10 media and filtered through a sterile 70 μm mesh. Splenic mononuclear cells were isolated by density gradient centrifugation using Ficoll-Paque. Viable cell suspensions from LN and spleen were cryopreserved in FBS + 10% DMSO and stored at -150°C until thawing.

### Flow cytometry acquisition and analysis

All antibody staining was done on freshly isolated cells following a protocol detailed elsewhere^92^. After washing with phosphate-buffered saline (PBS), cells (whole blood derived leukocytes or isolated lymphocytes) were prestained for the chemokine receptor CCR7 for 15 min at 37°C 5% CO_2_. All following incubations were performed at room temperature. Cells were stained for viability exclusion using Live/Dead Fixable Aqua (Invitrogen) for 10 minutes, followed by a 20-minute incubation with a panel of directly conjugated antibodies and Trustain (BioLegend) diluted with fluorescence-activated cell sorting (FACS) buffer (PBS containing 0.1% sodium azide and 1% bovine serum albumin) and Brilliant Stain Buffer (BD Biosciences). The cells were washed in FACS buffer and fixed in PBS containing 4% paraformaldehyde (Electron Microscopy Sciences). Cells were stored at 4°C in the dark until acquisition. All flow cytometry data was collected on a BD FACSymphony A5 cytometer (BD Biosciences). FlowJo software version 10.9 was used to generate population frequencies and representative flow cytometry plots. R was used for further analysis of these exported population frequencies.

### CITEseq sample processing and library generation

Eight samples were run simultaneously for each round of CITEseq, balancing both tissue type and disease state across runs. Cryopreserved cells were thawed by gentle shaking in a 37°C water bath until the cell suspension was partially thawed, which was immediately decanted into cold R10 medium + DNAse. Thawed cells were centrifuged at 500 xg for 7 minutes, then resuspended in room temperature R10 + DNAse. Cells were rested in R10 + DNAse at 2 x10^6^ cells/mL for 2 hr at 37°C with 5% CO_2_. All following cell preparation and antibody staining steps, including centrifugation, were done at 4°C. After resting, cells were centrifuged at 500 xg for 7 minutes and resuspended in 1 mL ice cold Cell Stain Buffer (CSB, BioLegend). Samples were filtered through a sterile pre-wet 70 μm mesh, followed by a 2 mL wash with CSB. 500k live cells that passed through the mesh were spun at 200 xg for 10 minutes and resuspended in CSB. Trustain was added to each sample at a 1:10 dilution and incubated on ice for 10 minutes. After, each sample received a unique hashing antibody (BioLegend) following manufacturer’s instructions. Totalseq-A antibody cocktail (BioLegend), reconstituted in CSB following the manufacturer’s protocol, was added to each sample at the manufacturer’s recommended concentration. After a 30 minute incubation on ice, cells were washed 3X with ice cold CSB, counted, and pooled in CSB at 2000 cells/μL with ∼125K cells from each sample. The pooled sample was filtered through a 40μm FlowMi strainer (Sigma) and counted. Pooled samples were run at 17.5k cells/well on a Chromium X (10X Genomics) using the Chromium Next GEM Single Cell 3’ HT or RT kit v3.1 (10X Genomics). Single cell sequencing library preparations for the RNA modality followed the manufacturer’s protocol, with the exception of spiking in primers at step 2.2 that amplify ADT and HTO sequences. 1 μL of 0.2 μM ADT additive primer (sequence: CCTTGGCACCCGAGAATT*C*C) and 1 μL of 0.1 uΜ ΗΤΟ additive primer (sequence: GTGACTGGAGTTCAGACGTGTGCTCTTCCGAT*C*T) was spiked into each individual sample prep. Preparation of the ADT and HTO single cell sequencing libraries followed manufacturer’s protocol (BioLegend) and described elsewhere (cite-seq.com)^93,94^. Briefly, at step 2.3d, 60 μL of supernatant was collected, 140 μL of SPRIselect reagent (Beckman Coulter) was added to the collected supernatant, washed twice with 80% ethanol, and eluted with 90 μL EB (Qiagen). For the respective ADT and HTO modalities, antibody-derived oligomers were amplified and indexed by mixing 45 μL eluent with 50 μL of KAPA HiFi HotStart ReadyMix (KAPA Biosystems) and 5 μL of forward/reverse indexing primers (Biolegend). Amplification settings for the ADT modality were as follows: 95 °C 3 min, ∼8 cycles of 95 °C 20 sec, 60 °C 30 sec, and 72 °C 20 sec, followed by 72 °C for 5 min and ending with hold at 4 °C. Amplification settings for the HTO modality were the same except for a 64 °C annealing temperature. Post-amplification, ADT and HTO libraries were purified with a 1.2X SPRIselect cleanup and eluted with 30 μL EB. All final libraries were quantified using a Qubit dsDNA HS Assay kit (Invitrogen) and a High Sensitivity D1000 DNA tape (Agilent) on a Tapestation D4200 (Agilent).

### CITEseq library sequencing

Sequencing runs were performed on the NovaSeq 6000 platforms (Illumina) with a target of at least 10,000 reads per cell for ADT libraries, 25,000 reads per cell for RNA libraries, and 500 reads per cell for HTO libraries.

### CITEseq preprocessing

The cellranger (7.0.0; 10X Genomics) suite of tools were used for all preprocessing steps for the RNA component. Raw BCL files were demultiplexed using the cellranger mkfastq tool, resulting in fastq files for each lane used in the 10X Genomics chip. Fastq files were processed into cell by gene matrices using the cellranger count tool by aligning against the hg38 reference genome. The resultant matrices were used for downstream analyses. For ADT and HTO, raw BCL files were demultiplexed using bcl2fastq2 (Illumina) to create fastq files that correspond to each lane used in the 10X Genomics chip. Fastq files were then counted using a variation of the kite pipeline (https://github.com/pachterlab/kite) which uses kallisto and bustools^95^ for alignment, barcode correction, and counting to create cell by feature matrices. Code for ADT and HTO preprocessing is found here: https://github.com/betts-lab/scc-proc.

### CITEseq modality processing and dehashing

The RNA filtered feature barcode matrix (as outputted from cellranger count) was loaded into R (v4.1.1) and Seurat (v4.1.1) to create a Seurat object after filtering out for wells that had low cell recovery and/or poor quality control metrics as determined by cellranger. Cells were then further filtered by the following criteria for initial quality checking: number of features > 200 & number of features < 6000 & percentage of mitochondrial reads < 12.5%. RNA counts were then normalized on a per cell basis. The ADT barcode matrix (as outputted from kallisto bustools) was loaded into the emptyDrops function (DropletUtils v1.14.2) with a minimum threshold of 100 UMI counts to determine what barcodes are associated with empty droplets (false discovery rate < 0.01). The empty droplets were filtered out of the ADT matrix. The intersection of usable barcodes was determined between the QC filtered RNA modality, the QC filtered ADT modality, and the HTO barcode matrix. The Seurat object was filtered to retain only barcodes that were detected/passing QC in all three modalities. After adding the ADT and HTO modalities as separate Assay objects, we normalized the HTO data within each hashtag. The HTODemux() function in Seurat with the argument (positive.quantile = 0.99) was used to demultiplex samples based on hashtag. Samples that were classified as singlets were kept for downstream analyses.

#### Modality integration and clustering

All runs were merged into a single Seurat object. ADT data were scaled and normalized using the centered-log ratio method by cell. Principal component analysis was performed for RNA and ADT separately and resultant principal components were each used as input for batch effect correction using Harmony^96^ with grouping variables set by donor and by each chip used for droplet emulsions. The batch effect corrected RNA-based principal components were used to generate a shared nearest neighbor (sNN) graph. Using the sNN graph, clustering was performed with the Leiden algorithm and the following settings for the FindClusters method in Seurat (method = “igraph” and resolution = 1). An uniform manifold approximation and projection (UMAP) representation of the data was constructed from the sNN graph for visualization purposes. Using both the ADT and RNA modalities, clusters were manually annotated. Clusters with multiplet signatures or phenotypes indicative of poor sample quality/viability were discarded. Given the known viability difficulties with human biopsy samples, we used a heat shock protein (HSP) gene set list (Group 582) from the HUGO Gene Nomenclature Committee (HGNC)^97^. These genes were fed into the AddModuleScore function in Seurat to generate a heat shock module. To remove confounding effects associated with heat shock response, cells with a score less than the 95th percentile of overall heat shock module scores were kept for downstream analysis.

### Weighted gene co-expression network analysis (WGCNA) for pancreatic lymph node samples

The dataset after clustering and quality control filtering were filtered for cells from the pancreatic lymph nodes and from donors associated with ND or T1D disease status for a more targeted analysis without confounding effects from other tissue sites and the inherent diversity of donors labeled as autoantibody positive. A WGCNA was constructed using the WGCNA^98^ and hdWGCNA^99^ packages where metacells were formed based on the following grouping variables: donor, disease status, and condensed phenotype from manual annotations. The resulting topological co-expression network was used to identify modules and assess correlations with disease status and manual phenotypes. All specific function parameters and steps for WGCNA are detailed in the study GitHub repository (see Resource Availability).

### HLA-GRS calculation and correlation to immune populations

Phasing of HLA-DQA1 and HLA-DQB1 genotypes was inferred via comparison with published haplotype frequencies from European Americans^100^. HLA-DQA1-DQB1 haplotypes were used to calculate the HLA component of the T1D polygenic risk score, GRS2, according to methods developed by Sharp, et al^68^. Scoring considered additive odds ratios for 14 individual haplotypes and interactions for 18 non-additive haplotype combinations as in the Polygenic Risk Score (PRS) Toolkit for HLA (v0.22a)^68^. Correlations between HLA-GRS and immune population frequency were computed using partial correlation analysis from the ppcor package.

### Graphics

All figures were made in R with the following packages: grid, ggplot2^101^, ComplexHeatmap, Seurat, hdWGCNA, and patchwork. All code to produce figures can be found in the study GitHub repository (see Resource Availability). Annotations were added to the R-generated figures using Adobe Illustrator (v.22.2.3).

### Statistical Analyses

For flow cytometry, all statistical analyses were run in R with the rstatix, ggpubr, and multicomp packages. Non-parametric analyses between two groups were performed with a two sample Wilcoxon test, with multiple hypothesis correction using Holm’s method when appropriate, using the rstatix package. Non-parametric analyses between 3 or more groups was performed with Dunn’s test, with Holm’s method for multiple hypothesis correction adjustment, using the rstatix package. Parametric comparisons between 3 or more groups were performed with robust one-way ANOVA in the WRS2 package, with post hoc testing using Hochberg’s multiple comparison adjustment and tr = 0.1. If cold ischemia was found to affect the frequency of an immune population, ANCOVA was used to control for the effect of cold ischemia on immune population frequency while comparing 3 or more groups, with post hoc analysis performed with Tukey’s test, using the multicomp package. Each dot in a plot depicting summarized flow cytometry data is an individual sample. Sample distributions represented by a boxplot, with the median depicted as the center value, quartile ranges forming the box edges, and whiskers depicting the minimum to maximum distribution. Correlations between HLA-GRS and immune population frequency were run using the partial Kendall rank correlation while controlling for the confounding variables of disease state and cold ischemia time. Correlation p-values were adjusted for multiple comparisons using Benjamini-Hochberg multiple test correction. For CITEseq analysis, samples were quality controlled as explained in the “CITEseq modality processing and dehashing” section. To help elucidate more specific cell type differences in differential gene analysis, a common set of differentially expressed genes was computed for each manually annotated cluster between disease states using the Wilcoxon Rank Sum implementation in Seurat FindAllMarkers with a log fold change threshold of 0.1. Any differential genes that were found in more than 12 clusters were labeled as common genes that were largely independent of cell phenotype. Combinatorial differential expression testing was performed with Seurat FindMarkers (Wilcoxon Rank Sum) followed by Bonferroni correction for multiple tests calculated across all tests performed across the combinations. Unless otherwise stated in text or figure legends, significance by statistical test was set as adjusted p value < 0.05. * is p < 0.05, ** is p < 0.01, *** is p < 0.001 for all figures. Refer to figure legends for the application and details of each test.

## Data Availability

All de-identified human flow cytometry, sequencing-based, clinical data have been deposited on PANC-DB (https://hpap.pmacs.upenn.edu/). Raw sequencing -based data are also deposited in Genbank under accession number GSE221787. All original code has been deposited at GitHub at https://github.com/betts-lab/hpap-tissue-citeseq. Exceptions are the code used to generate HLA-GRS, which can be found at https://github.com/sethsh7/hla-prs-toolkit version 0.22a^68^, and code for ADT and HTO preprocessing, which is found at https://github.com/betts-lab/scc-proc. All data are publicly available as of the date of publication. Additional data from histology, immunophenotyping, repertoire profiling, metabolic and transcriptomic studies are generated on all HPAP donors and are deposited on PANC-DB (https://hpap.pmacs.upenn.edu/), providing a shared and publicly available comprehensive resource.

## Code Availability

All original code has been deposited on GitHub at https://github.com/betts-lab/hpap-tissue-citeseq. Exceptions are the code used to generate HLA-GRS, which can be found at https://github.com/sethsh7/hla-prs-toolkit version 0.22a^68^, and code for ADT and HTO preprocessing, which is found at https://github.com/betts-lab/scc-proc. All code are publicly available as of the date of publication.

