## Supplemental Files for "Immune perturbations in human pancreas lymphatic tissues prior to and after type 1 diabetes onset"

### **Supplemental Information**

Extended Data Figure Legends and Extended Data Tables 1-2

### **Supplemental Legends**

**Extended Data Figure 1. Gating strategy for flow cytometry**

Representative gating strategy of HPAP-099 splenocytes for the lineage flow cytometry panel. PBMCs = peripheral blood mononuclear cells, NK = natural killer, DCs = dendritic cells, ILCs = innate lymphoid cells, HSCs = hematopoietic stem cells, Tcm = central memory T cell, Tem = effector memory T cell, Temra = effector memory CD45RA+ T cell, Tregs = regulatory T cells, gcTfh = germinal center follicular helper T cell, mTfh = mantle follicular helper T cell, Tn = naive T cell, Tnl = naive-like T cells (all non CD27+CD127+ Tnaive)

**Extended Data Figure 2. Distribution of immune populations across tissues**

(A) NK cells, (B) B cells, (C) CD Tn, (D) T cells, (E) ILC, and (F) CD8 Tcm distribution across tissues. * is p < 0.05, ** is p < 0.01, *** is p < 0.001, as determined by robust one-way ANOVA, with post hoc testing using Hochberg’s multiple comparison adjustment.

(G) PCA of major immune lineage populations and (H) CD4+ T cell subsets in the lymph nodes.

(I) Hierarchical clustering of all lymph node samples within the flow cytometry dataset using CD4+ T cell subset populations. Coloring of the heatmap represents an individual sample’s Z-score within the respective immune population.

**Extended Data Figure 3. CITEseq cluster phenotype**

(A) Surface antibody and (B) RNA expression of a subset of genes used to annotate the CITEseq dataset. Average expression is scaled on a per-gene basis.

**Extended Data Figure 4. WGCNA modules between ND and T1D donors**

A) All other WGCNA modules not present in Figure 3. Modules were generated using WGCNA analysis on scRNAseq data from pLN lymphocytes in ND and T1D disease states.

(B) Correlations of modules in (A) between ND and T1D disease states, across all immune cell clusters . * is p < 0.05, ** is p < 0.01, *** is p < 0.001.

**Extended Data Figure 5. Immune cell phenotypes in the mLN and spleen**

(A) Frequency of CD25+ CD4+ T cells, as detected by flow cytometry, in mLN and (B) spleen. Statistical significance determined by robust ANOVA with post hoc testing using Hochberg’s multiple comparison adjustment. Boxplot represents median and interquartile range.

(C) Frequency of CD25+ CD127- CD4+ Tregs, as detected by flow cytometry, in mLN and (D) spleen. Statistical and graphical information the same as above.

(E) Expression of CD4+ Treg-associated genes within the CD4+ Treg/Tcm cluster from mLN and (F) spleen. Expression scaled within each gene., as determined by Wilcoxon Rank Sum Test and p-value adjustment using the Bonferroni method.

(G) Frequency of CD4+ Tn, as detected by flow cytometry, in mLN and (H) spleen. Statistical and graphical information the same as in (A).

(I) Frequency of CD25+ CD8+ T cells in mLN and (J) spleen, as detected by flow cytometry. Statistical and graphical information is the same as in (A).

(K) Frequency of CD38+ CD8+ T cells in mLN and (L) spleen, as detected by flow cytometry. Statistical and graphical information is the same as in (A).

(M) Normalized expression of *CXCR3* and *TOX* in CD8 Tcm/Tem/Temra cells in the mLN and (N) spleen. P-value determined by Wilcoxon Rank Sum test with Bonferroni correction done between all combinatorial tests. Dashed red line indicates the exclusive threshold (> 0.5) for demarcating positive gene expression. Gray arrows indicate direction of expression relative to ND.

(O) Normalized expression of *GZMB* and *KLRB1* within the combined NK and NK/ILC clusters in the mLN and spleen. Statistical significance tested using Wilcoxon Rank Sum Test with p-value adjustment using the Bonferroni method.

For all panels in this figure, * is p < 0.05, ** is p < 0.01, *** is p < 0.001

**Extended Data Figure 6. Phenotype of naive T cell clusters in the CITEseq dataset**

(A) Surface protein expression and (B) RNA expression of a set of genes used to delineate naive T cell clusters and memory T cell clusters.

**Extended Data Table 1**

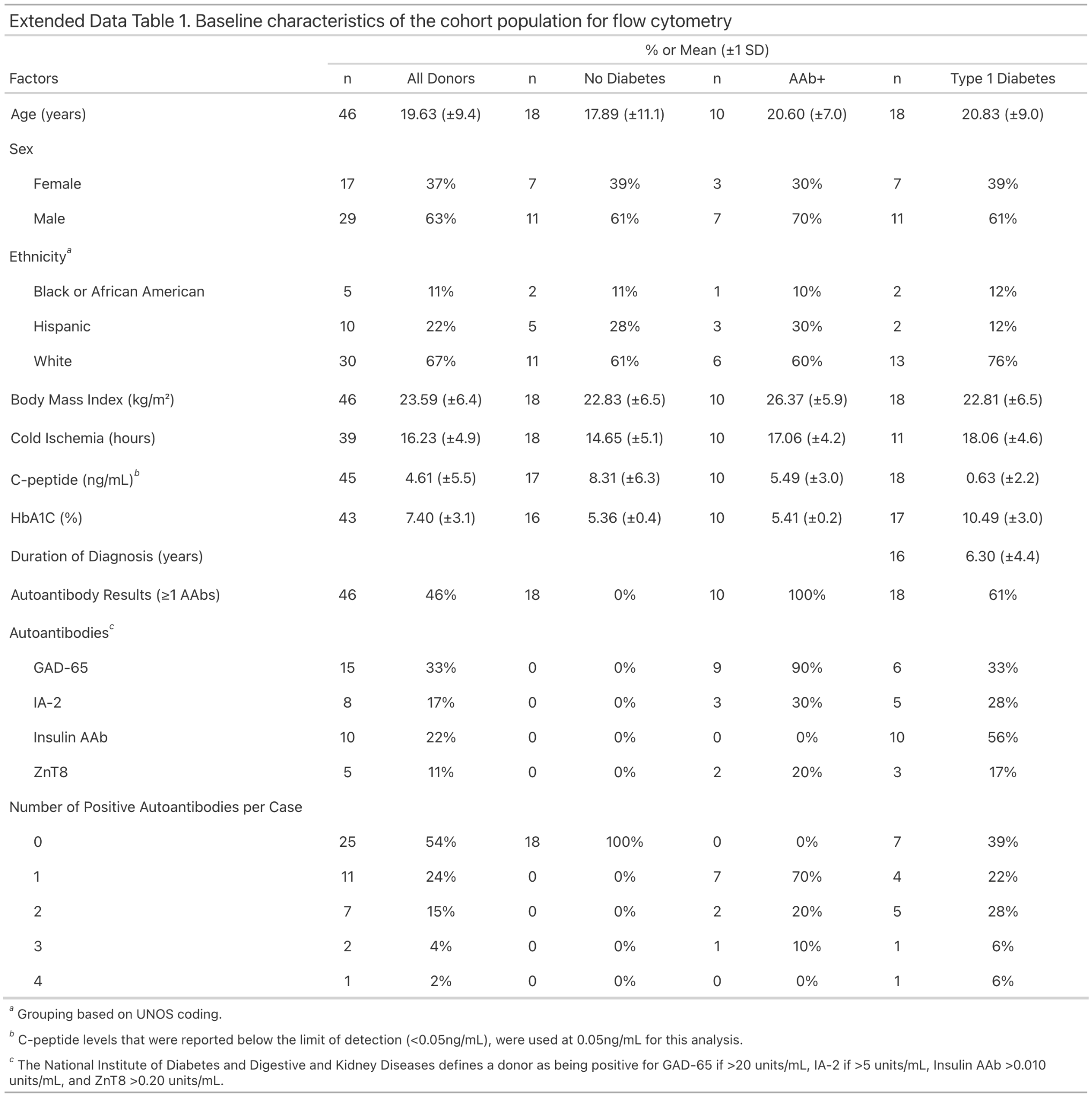

### **Extended Data Table 2**

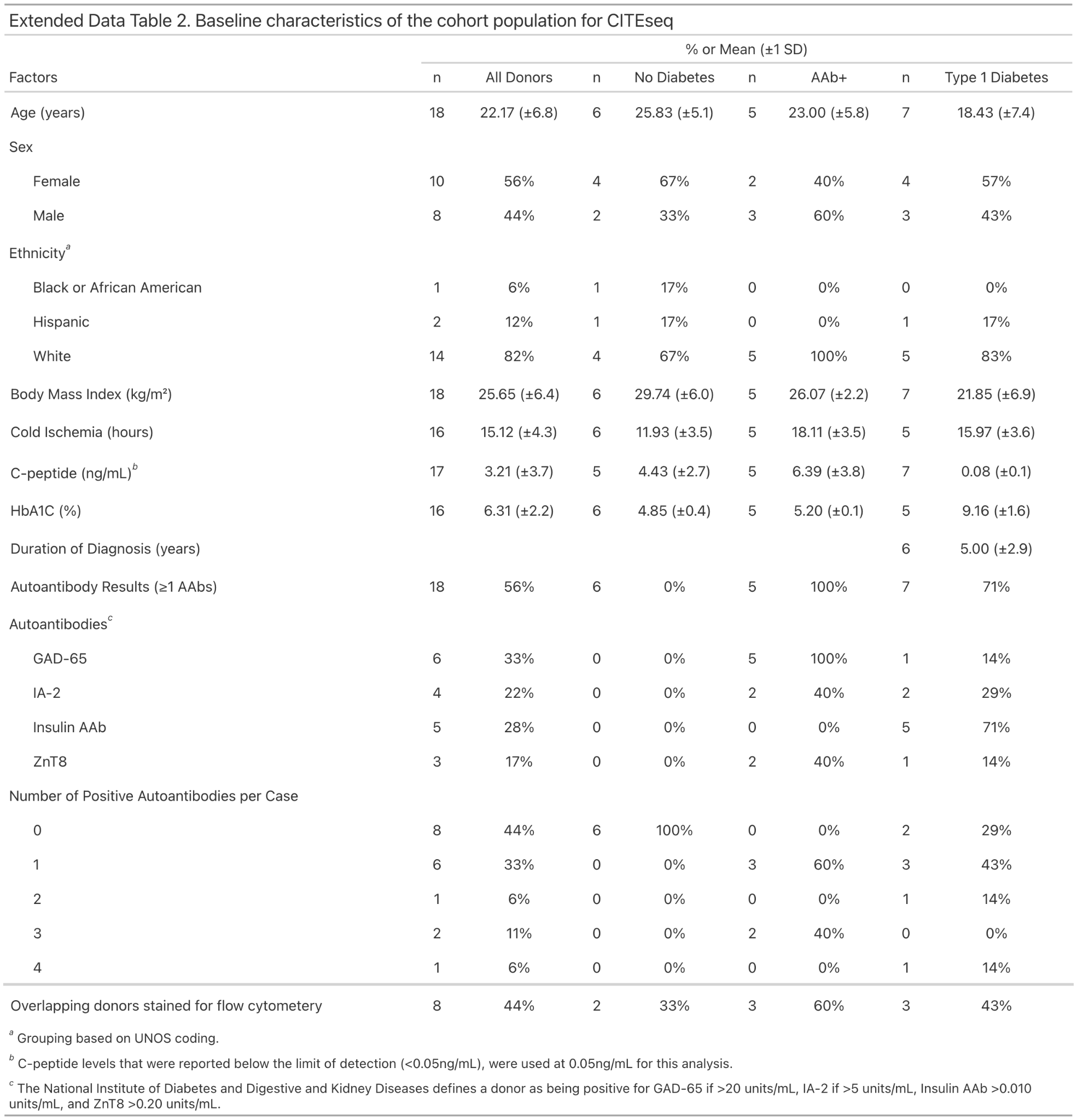
