## Supplementary figures and images for "Immune perturbations in human pancreas lymphatic tissues prior to and after type 1 diabetes onset"

### Supplemental Figure 1

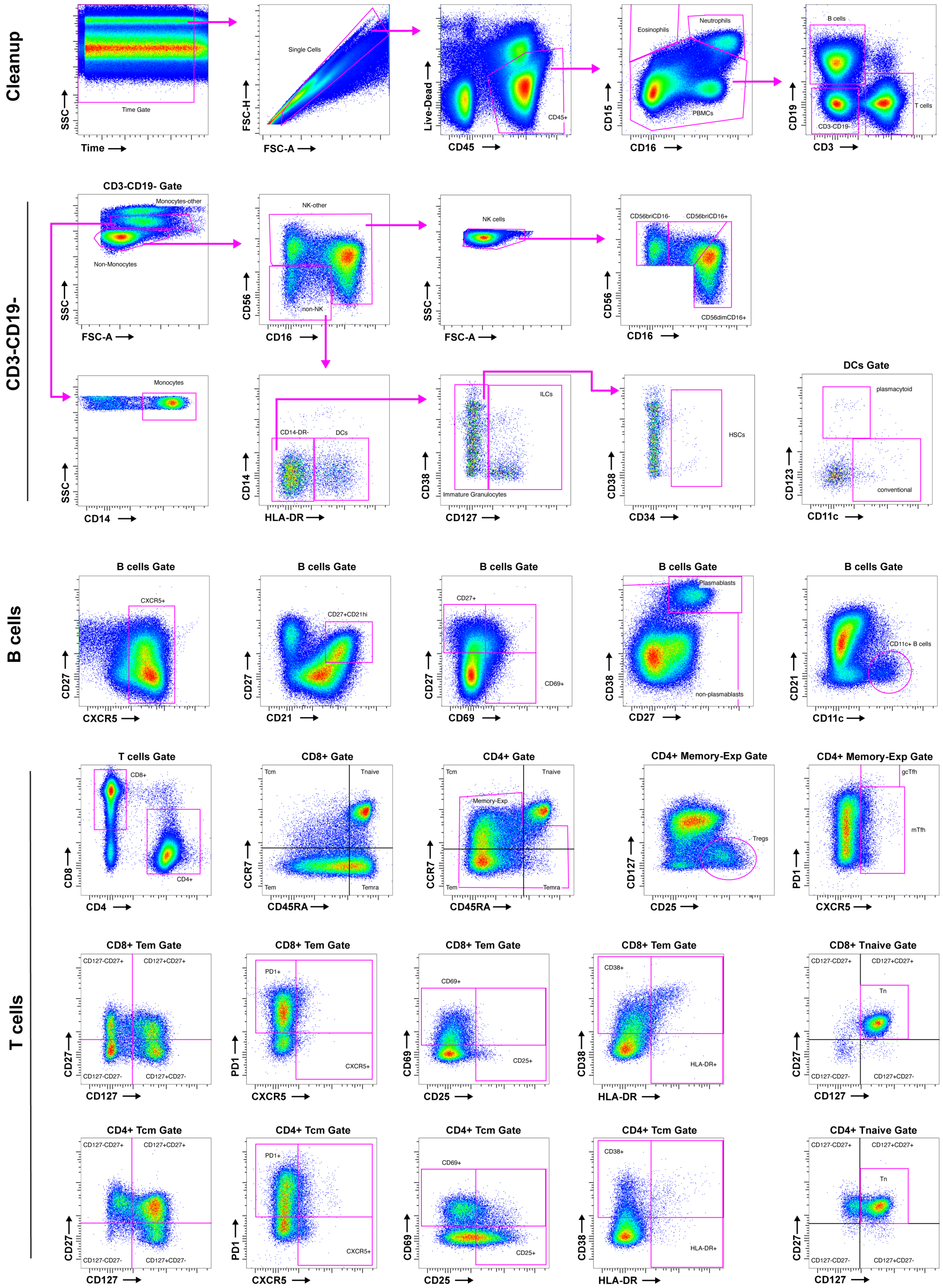

### Supplemental Figure 2

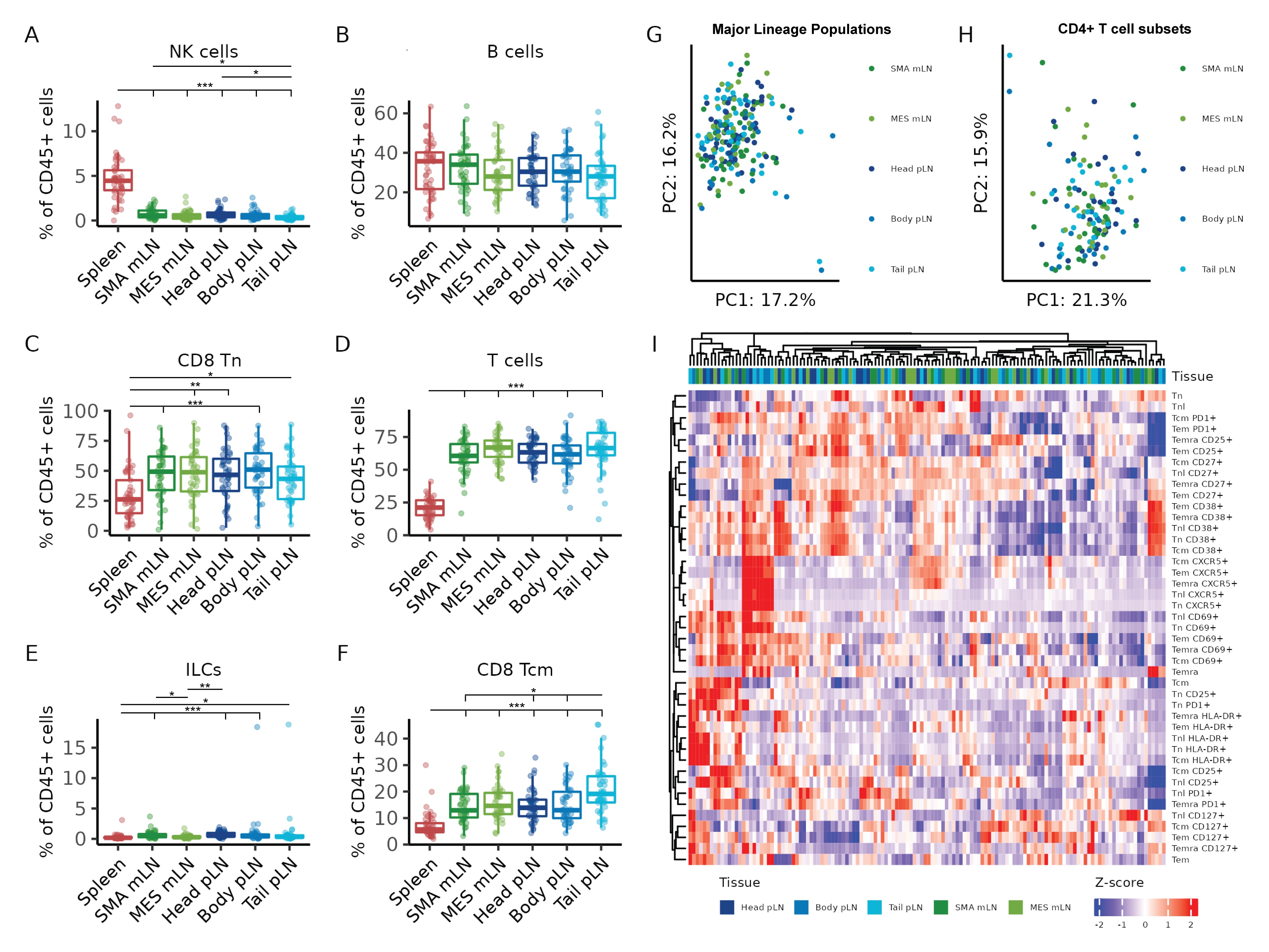

### Supplemental Figure 3

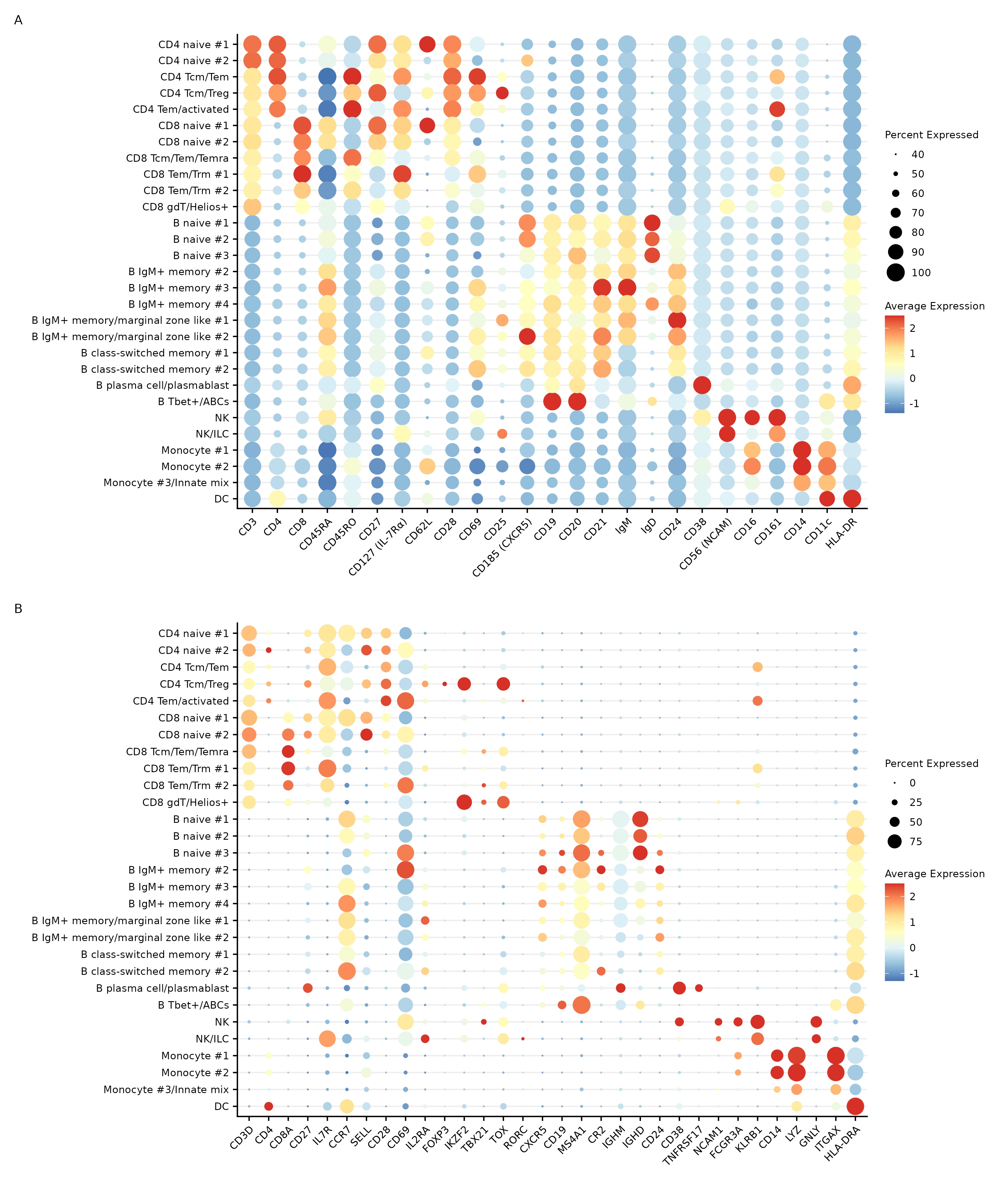

### Supplemental Figure 4

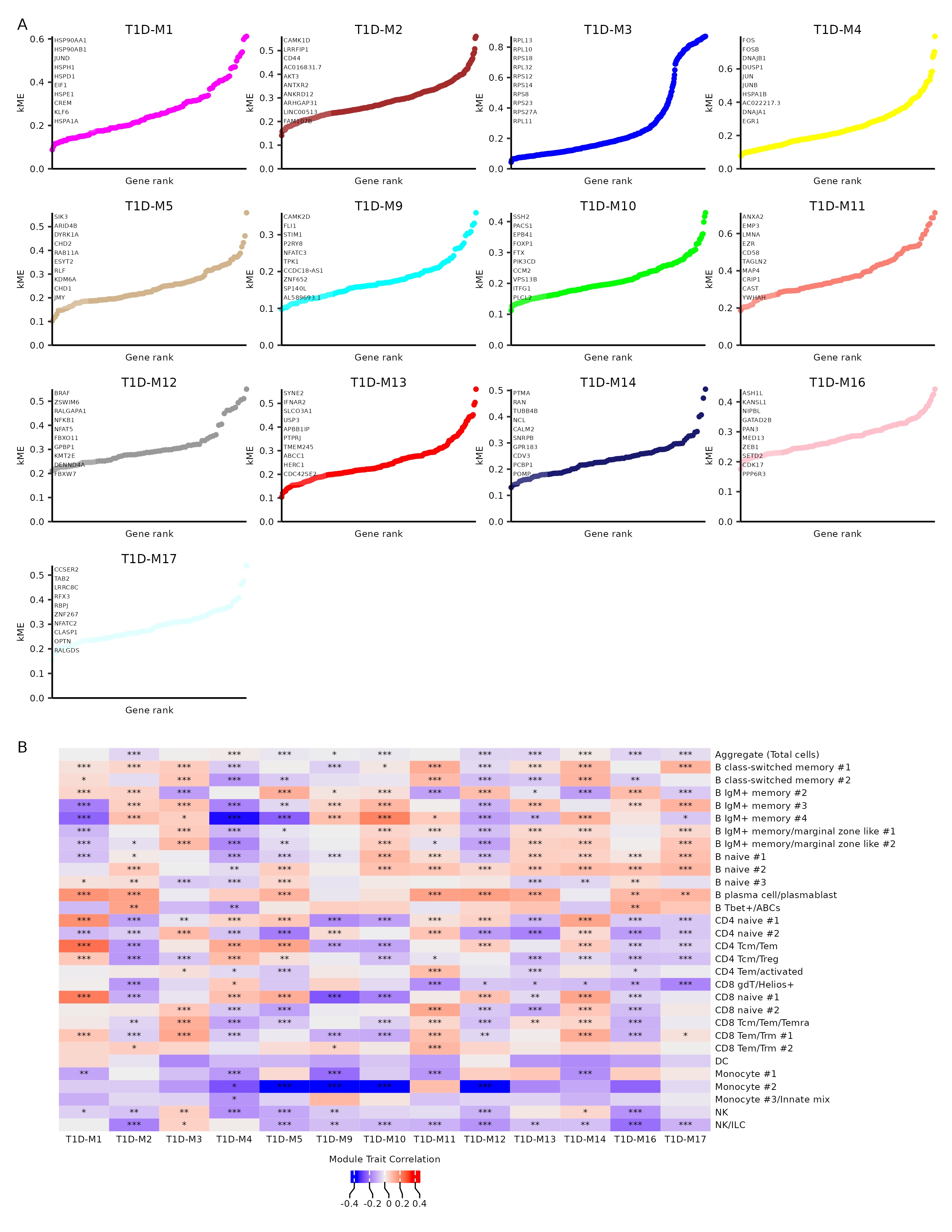

### Supplemental Figure 5

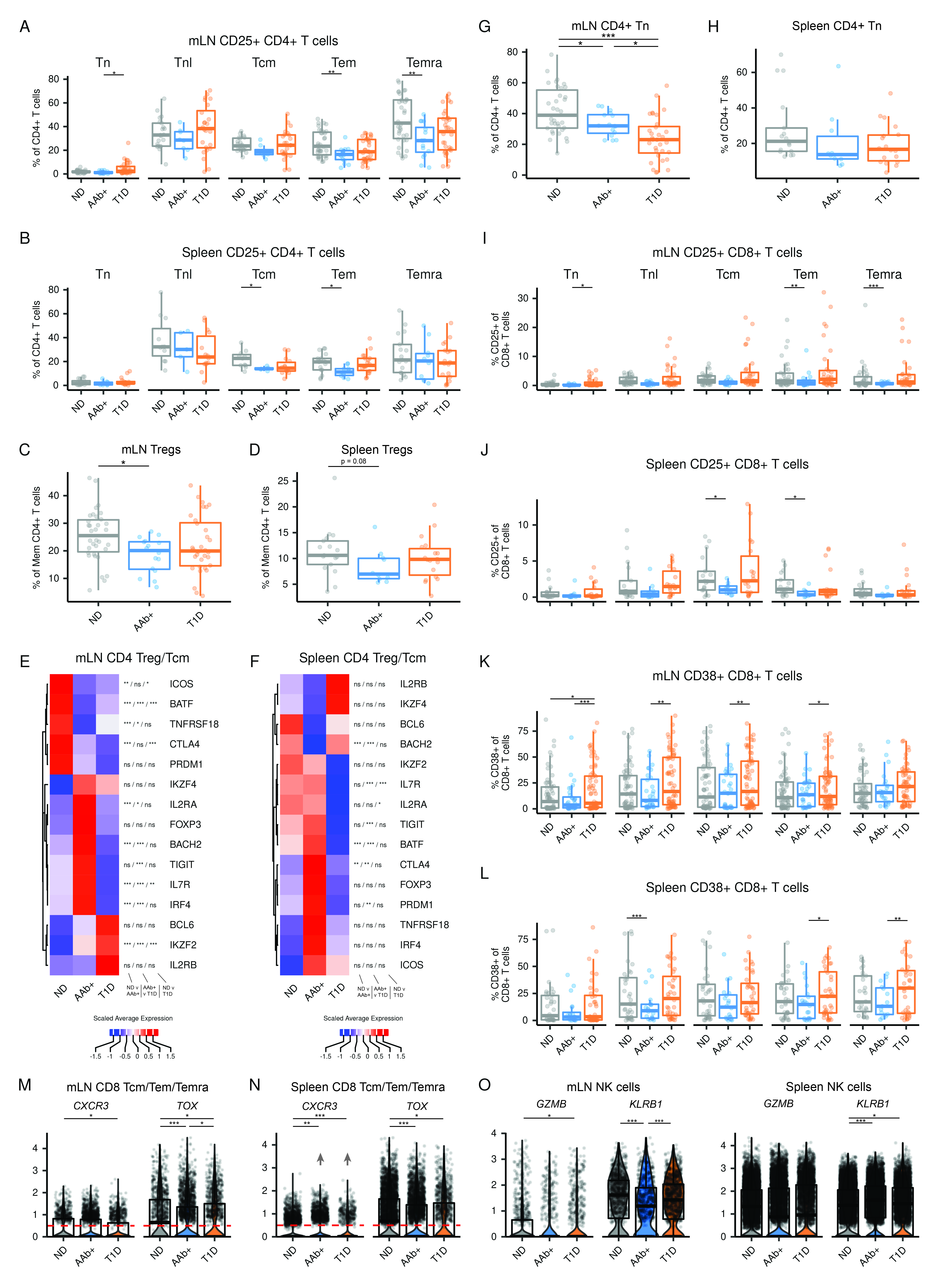

### Supplemental Figure 6

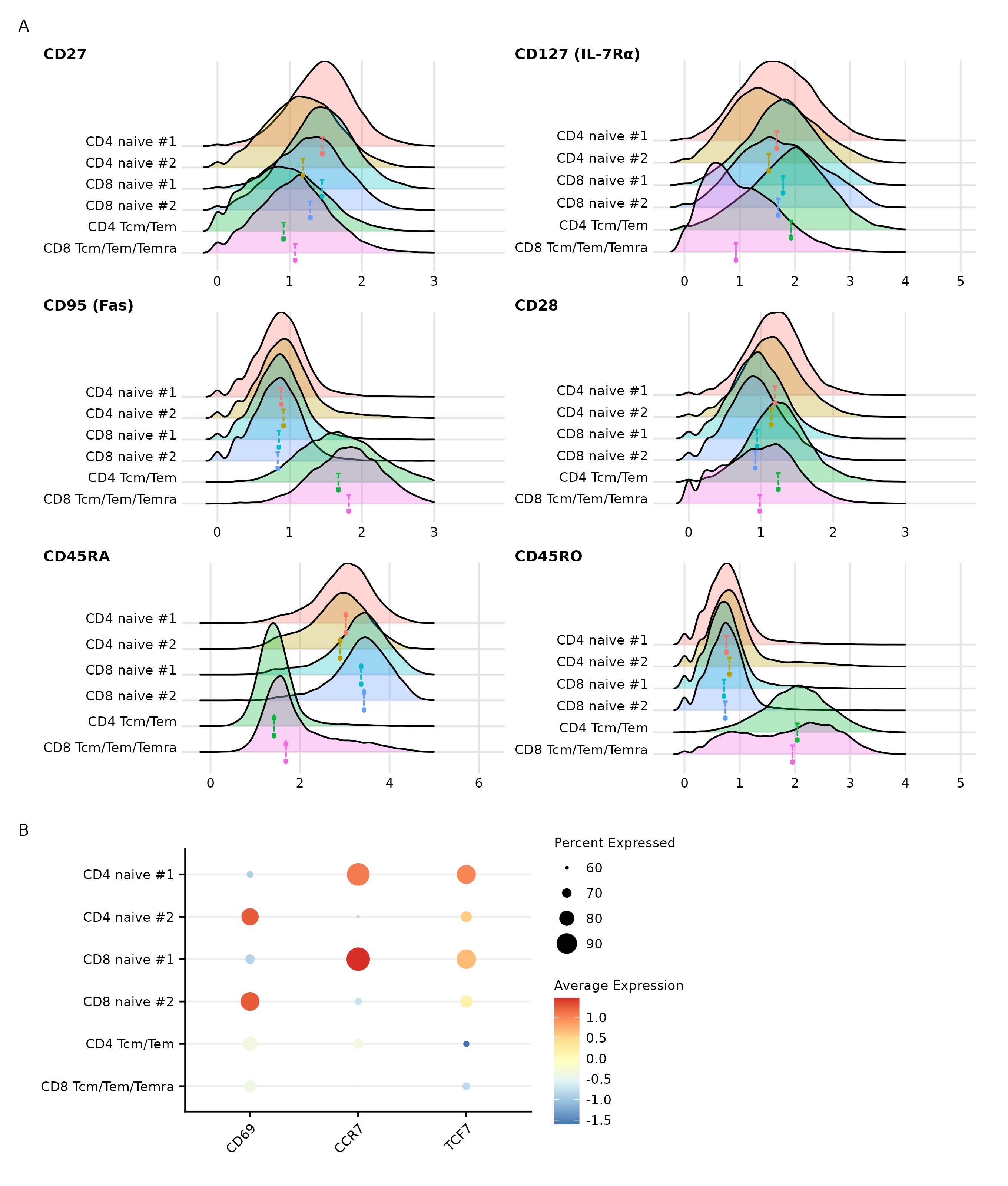
